# Manipulating vector transmission reveals local processes in bacterial communities of bats

**DOI:** 10.1101/2021.03.03.433743

**Authors:** Clifton D. McKee, Colleen T. Webb, Michael Y. Kosoy, Ying Bai, Lynn M. Osikowicz, Richard Suu-Ire, Yaa Ntiamoa-Baidu, Andrew A. Cunningham, James L. N. Wood, David T. S. Hayman

## Abstract

Infectious diseases result from multiple interactions among microbes and hosts, but community ecology approaches are rarely applied. Manipulation of vector populations provides a unique opportunity to test the importance of vectors in infection cycles while also observing changes in pathogen community diversity and species interactions. Yet for many vector-borne infections in wildlife, a biological vector has not been experimentally verified and few manipulative studies have been performed. Using a captive colony of fruit bats in Ghana, we observed changes in the community of *Bartonella* bacteria over time after the decline and subsequent reintroduction of bat flies. With reduced transmission, community changes were attributed to ecological drift and potential selection through interspecies competition mediated by host immunity. This work demonstrated that forces maintaining diversity in communities of free-living macroorganisms act in similar ways in communities of symbiotic microorganisms, both within and among hosts. Additionally, this study is the first to experimentally test the role of bat flies as vectors of *Bartonella* species.

## Introduction

Knowledge of the processes driving parasite diversity is central to understanding infection dynamics in endemic populations and pathogen emergence in new hosts. In contrast to an historical focus on simple one-host, one-parasite systems, there is now greater appreciation that parasites exist within communities of other parasites, harbored by hosts that may vary in their responses to parasitism (Johnson et al., 2015). Yet it is not clear how well ecological theory developed for free-living organisms applies to communities of microorganisms (Sutherland et al., 2013). This is especially true for parasites and symbionts due to the environmental feedbacks that exist from their dependence on hosts for survival and reproduction (Costello et al., 2012; Miller et al., 2018). Additionally, parasite community dynamics within hosts may occur at differing timescales compared to transmission among hosts. Given these differences, experimental manipulations of natural parasite communities are needed to explore the generality of community theory across organisms.

The metacommunity concept is a useful framework to apply toward analyzing parasite community dynamics within hosts (Leibold et al., 2004; Mihaljevic, 2012). In this framework, hosts are discrete patches harboring potentially interacting parasite species. Similar to free-living organisms, four forces might be expected to affect parasite community diversity: speciation, dispersal, ecological drift, and ecological selection (Vellend, 2010). Within a metacommunity, the relative importance of these forces may vary at different scales (Seabloom et al., 2015), i.e., within versus among hosts. Speciation is the only force that generates parasite diversity *de novo*, but is generally slow and dependent upon dispersal for newly created diversity to penetrate to all scales. Dispersal is the movement of parasite species within a host, among hosts through transmission, or among host populations through host movement. Within metacommunities, parasite species with equal competitive ability may vary stochastically in the production of new parasite individuals or in new infections through transmission. This ecological drift can lead to changes in community composition within hosts (e.g., loss of rare species) or among hosts (e.g., increases in beta diversity), similar to predictions of neutral theory (Hubbell, 2001). Drift happens faster in small communities with few parasite individuals and with little dispersal. Lastly, ecological selection acts within and among hosts. Selection occurs because parasite species vary in replication success within different host individuals or species because of variation in susceptibility or tolerance. Additionally, parasite species may compete within a host, either indirectly through shared resources or common enemies, such as the host immune system, or directly through interference (Pedersen & Fenton, 2007). Species with higher success within a host will dominate and may exclude others, but this can be counterbalanced if fitness is driven by dispersal ability over interspecific competition or there is frequency-dependent selection by the host immune system. These four forces could separately affect parasite community diversity over time. While speciation ultimately creates diversity, the other forces sort parasite species across scales. Thus, a strategy for studying parasite community diversity is to understand the relative importance of these forces both within and among hosts (Seabloom et al., 2015).

Manipulative experiments are one approach to measuring the relative influence of ecological forces acting on communities. By changing the strength of one force, one can observe how others respond and interact across scales. While previous studies have performed parasite community manipulations within and among hosts (see Mihaljevic, 2012 and Johnson et al., 2015 for examples), few studies to our knowledge have looked at how manipulating forces that act across scales lead to changes in other forces. Since dispersal is the force that interacts with other processes across within-host and among-host scales (Vellend, 2010), it is an appealing target for manipulation.

Vector-borne infections are ideal systems for experimental study because reduction of vector density limits dispersal of parasites between hosts, allowing the analysis of other forces affecting the relative abundance of parasite species. Using a captive colony of straw-colored fruit bats (*Eidolon helvum*) in Ghana, the community dynamics of *Bartonella* bacteria were monitored in bats over three years. During this experiment, the presumed vectors (bat flies) declined in density within the colony but were then reintroduced. The experiment thus controls parasite dispersal across two scales: the captive colony is closed to immigration (pups enter the colony uninfected) and transmission is manipulated via changes in the bat fly population size. By manipulating parasite dispersal, the effect of among-host dispersal is minimized and the effects of local, within-host effects (ecological drift and selection) on parasite dynamics and diversity can be observed. We hypothesize that *Bartonella* communities in the colony will respond to changes in among-host dispersal/transmission by bat flies. Specifically, we predict that infection prevalence and diversity will at first decline concurrently with the bat fly population and then increase upon reintroduction of flies, thus providing experimental evidence that bat flies are vectors of *Bartonella* in bats. We hypothesize that limitation of parasite dispersal will result in stochastic losses of rare *Bartonella* species and increases in community beta diversity due to ecological drift, and shifts in the rank abundance of *Bartonella* communities due to local selection. Finally, potential interactions among *Bartonella* species will be detectable based on coinfection frequencies, specifically evidence of competition and/or facilitation. This work expands our understanding of *Bartonella* dynamics in natural communities, particularly in bats and their ectoparasites. More broadly, this experiment deepens our understanding of the processes that affect parasite communities, patterns which may be compared with those seen in communities of free-living or mutualistic organisms.

## Materials and methods

### Study system

*Eidolon helvum* (Chiroptera: Pteropodidae) is a long-lived, tree-roosting bat species that can form enormous colonies during the local dry season (Fahr et al., 2015; Hayman et al., 2012). Bat flies (*Cyclopodia greefi*; Diptera: Nycteribiidae) are obligate blood-feeding ectoparasites of bats. The flies are wingless but can move among hosts within densely populated roosts.

*Bartonella* spp. (Alphaproteobacteria: Rhizobiales) are intracellular bacteria that infect mammals and are transmitted by blood-feeding arthropods (Harms & Dehio, 2012). At least six distinct *Bartonella* species have been previously described in *E. helvum* (Bai et al., 2015; Kosoy et al., 2010) and the same species plus additional variants have been detected in *C. greefi* (Billeter et al., 2012; Kamani et al., 2014). Based on these data and other studies (Brook et al., 2015; Morse et al., 2012; Moskaluk et al., 2018), it has been proposed that bat flies are vectors of *Bartonella* spp. in bats, but no experimental studies have been performed to demonstrate their competence.

Materials for this study come from a captive population of *E. helvum* bats in Accra, Ghana (Baker et al., 2014). Briefly, the captive facility is a double-fenced hexagonal 27.5 m diameter and 3.5 m high structure; a solid metal roof and cladding at the base prevent contact with other animals. The captive population was founded by three cohorts (Table S1) of mixed age and sex (n = 78) collected from a large seasonal colony in Accra (Hayman et al., 2012). The cohorts entered the colony in July 2009, November 2009, and January 2010; two additional cohorts were born in captivity in April 2010 (produced by mating between wild bats before entering the colony) and 2011 (produced by mating in captivity). All 13 captive-born neonates were matched to the dam they were attached to at the first sampling point after birth. Ethics approval for bat capture and the fly reintroduction experiment was obtained from the Zoolgical Society of London Ethics Committee (WLE/0467), the Veterinary Services Directorate of Ghana, and the Wildlife Division of the Forestry Commission of Ghana.

Bats were assigned to age classes and sex upon entry to the colony and afterward according to approximate birth date and secondary sexual characteristics (Peel et al., 2016): neonate, juvenile, sexually immature adult, and sexually mature adult. Passive integrated transponder (PIT) tags were implanted in each bat either at entry or shortly after birth to uniquely identify each bat and adult bats additionally received necklaces with alphanumeric codes. Although 112 total bats entered the colony, 25 bats left the colony either through recorded mortality (n = 12) or presumed mortality after being recorded missing for ≥3 sampling points (n = 13). Furthermore, not all bats had complete sample histories throughout the experiment because they intermittently escaped capture for processing.

Blood samples were taken from the captive bats every two months in 2009 and 2010 and every four months in 2011 (Table S1; see Appendix 1 for sampling protocol). On 6 March 2010 (denoted M10, day 221), a sample of bat flies (*C. greefi*; n = 28) was removed from the colony for testing for the presence of *Bartonella* DNA and from that point forward the fly population was observed to decline. Subsequent to this, it is assumed that little among-host bacterial transmission was occurring. To test the effect of restoring transmission on *Bartonella* community dynamics and to provide evidence that bat flies are vectors, bat flies were experimentally reintroduced to the colony. On 17 January 2012 (J12, day 903), a sample of adult bat flies and nymphs was taken from the original wild source colony of bats (n = 51), along with matched blood samples from donor bats (n = 42), and the flies were randomly assigned to approximately half the bats in the colony (n = 40; 1–4 flies per bat) while additional bat flies were collected for testing for the presence of *Bartonella* DNA (n = 18). Blood samples from captive bats were subsequently taken at three additional time points after the reintroduction of flies: 31 January 2012, 14 February 2012, and 15 March 2012. In total, 910 blood samples were taken from the captive colony over 14 time points from 2009 to 2012 (a period of 961 days), of which 905 samples could be definitively assigned to an individual by PIT tag or necklace ID. An additional 50 blood samples and 18 flies were taken from wild bats on J12.

### Bacterial detection and gene sequencing

The focus of this study was on changes in *Bartonella* infection prevalence and the relative abundance of different *Bartonella* species in bats, so a molecular detection and sequencing approach capable of distinguishing coinfecting species was used. Bat blood and fly samples were tested for the presence of *Bartonella* DNA using a multi-locus PCR platform (Bai et al., 2016) targeting fragments of the 16S–23S ribosomal RNA intergenic spacer region (ITS), citrate synthase gene (*gltA*), and cell division protein gene (*ftsZ*). Each of these loci is capable of distinguishing among *Bartonella* species and subspecies (La Scola et al., 2003), but may have amplification biases toward different *Bartonella* species in a sample (Himsworth et al., 2020; Kosoy et al., 2018). Thus, the purpose of this multi-locus approach was to confirm the detection of *Bartonella* DNA and to indicate across loci whether infections with multiple species were present. Further quantification of *Bartonella* infection load was performed using real-time PCR targeting the transfer-messenger RNA (*ssrA*). Sequences were verified as *Bartonella* spp. using the Basic Local Alignment Search Tool (BLAST; https://blast.ncbi.nlm.nih.gov/Blast.cgi).

Samples were only considered positive if a significant match was observed, even if there was a positive real-time PCR result (cycle threshold value [Ct] < 40). *Bartonella* sequences with multiple peaks in the electropherogram were separated into two or more distinct sequences by comparison with previously obtained *Bartonella* sequences from *E. helvum* and *C. greefi* (Bai et al., 2015; Billeter et al., 2012). Due to the frequency of multiple sequences obtained from these loci, conflicting sequences across genes were interpreted as evidence of coinfection rather than homologous recombination, and thus we report counts of sequences from distinct *Bartonella* species within a sample as recommended by Kosoy et al. (2018). All variants of *Bartonella* sequences sharing <95% sequence similarity with previously identified *Bartonella* species were submitted to GenBank. Additional details on bacterial detection and phylogenetic analysis are provided in Appendix 1.

### Data recording and statistical analyses

Relevant measures of *Bartonella* infection prevalence, infection load, and diversity were recorded or calculated to assess changes that occurred during the experiment, including before and after the reintroduction of bat flies to the captive colony. *Bartonella* infection prevalence within the captive bat colony, in sampled wild and captive flies, and from wild bats was reported based on the number of tested bats or flies that were positive at one or more loci (ITS, *gltA*, *ftsZ*, *ssrA*). Wilson scores were used to calculate 95% confidence intervals for single infection and coinfection prevalence. *Bartonella* alpha diversity was measured by *Bartonella* species richness and Shannon number, i.e., the effective number of species or the exponent of Shannon’s diversity index (Jost, 2006); species richness within each sample based on the number of loci positive was also recorded. *Bartonella* species relative abundances were calculated from the total number of sequences obtained across all loci, including separate sequences obtained from the same locus. A custom bootstrapping procedure with 1000 samples from the observed multinomial distribution of *Bartonella* species relative abundances was used to estimate 95% confidence intervals around measures of alpha diversity. *Bartonella* beta diversity was measured across sampled bats and flies using the binomial index option of the vegdist function in the R package *vegan* (Oksanen et al., 2019; R Core Team, 2020). Infection load was recorded as the number of loci positive and real-time PCR Ct value for each sample. Additionally, for each bat the time until becoming infected after first entering the colony and the duration of infection for the most persistent *Bartonella* species were recorded. These measures help to track whether certain demographic groups are more affected by the reintroduction of flies and to compare with changes in relative abundances of *Bartonella* species over time, respectively. Change points in *Bartonella* prevalence, infection load, and diversity measures were detected with segmented regression using the R *segmented* package (Muggeo, 2020). Chi-square or Fisher’s exact tests were performed to compare changes in infection status for bats that did or did not receive bat flies on J12. Multinomial and binomial likelihood ratio (LR) tests adapted from Pepin et al. (2013) were performed to find statistical associations between coinfecting *Bartonella* species and to detect changes in the relative abundances of *Bartonella* species during the study period. For additional details regarding regression analyses and likelihood ratio tests, see Appendix 1.

## Results

### Phylogenetic analysis of detected bacteria

*Bartonella* infections in bats and bat flies were identified as six previously characterized species based on ITS, *gltA*, and *ftsZ* sequences: *Bartonella* spp. E1–E5 and Ew (Bai et al., 2015; Kosoy et al., 2010). Two additional genogroups identified by *gltA* sequences, *Bartonella* spp. Eh6 and Eh7 (Figure S1), were similar to sequences previously obtained from *C. greefi* collected from *E. helvum* in Ghana and two islands in the Gulf of Guinea (Billeter et al., 2012). Phylogenetic analysis of concatenated *ftsZ* and *gltA* sequences distinguished Eh6 and Eh7 from other *Bartonella* species associated with *E. helvum* or other bat species (Figure S4). See Appendix 2 for more details on phylogenetic analysis.

### Bartonella infection prevalence and effects of bat fly reintroduction

As predicted, *Bartonella* prevalence in the captive colony changed with the population density of bat flies. *Bartonella* prevalence in the first three cohorts was high at colony entry, then declined concurrently with the observed decline in the bat fly population (Figure 1A). After flies were reintroduced, prevalence increased from 31% on day 903 to 48% on day 961. This change is reflected in the segmented regression analysis (Figure S6A; Table S4) with a shift from positive to negative slope near M10 (day 221) and a shift from negative to positive slope near J12 (day 903). The trend in *Bartonella* prevalence in the colony over time was similar if bats were considered positive for *Bartonella* with a threshold of at least one, at least two, at least three, or all genetic markers being positive (Figure S7).

**Figure 1.**
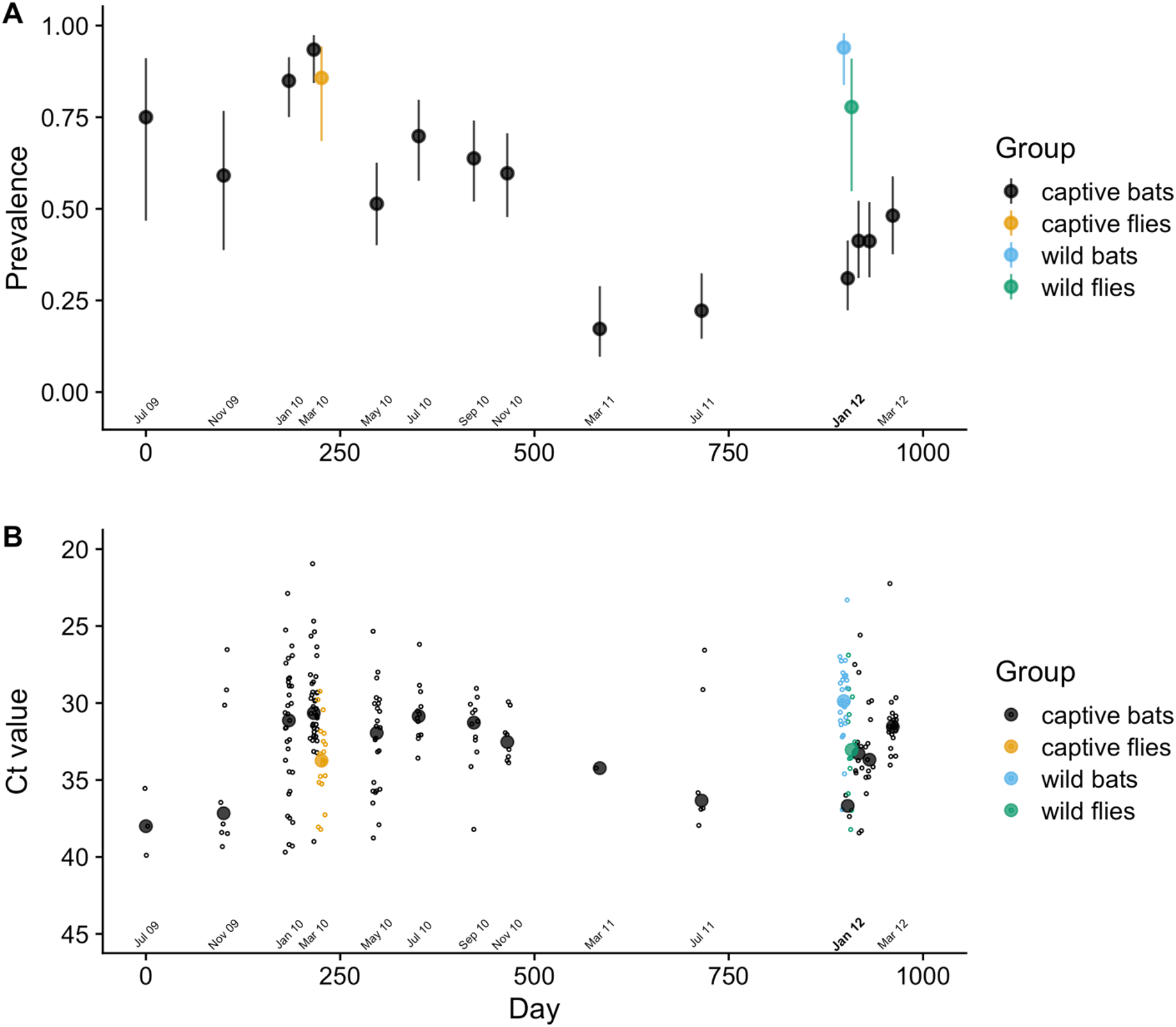
*Bartonella* infection prevalence and load in a captive colony of *E. helvum* over time. (A) Bats and bat flies were considered positive if a *Bartonella* sequence was obtained from one or more genetic markers. Wilson score 95% confidence intervals were drawn around prevalence estimates at each sampling time point. (B) Only points with RT-PCR Ct values < 40 are shown. Mean Ct values calculated at each time point are drawn as a filled circles over the data (open circles). The month labeled in bold font on the x-axis shows when bat flies were reintroduced.

The effect of bat fly reintroduction affected some age classes of bats more than others. Most sexually immature and sexually mature adult bats initially entered the colony infected (Figure S8A). All sexually immature bats were infected at entry and at the end of the study, but there was an increase in the proportion of sexually mature adult bats (χ^2^ = 3.2, df = 1, P = 0.038) infected by the end of the study compared to the start. Bats born into the colony in 2010 and 2011 were *Bartonella*-negative at first sampling. By the end of the experiment, 88% of these bats had become infected (Figure S8A), a very significant increase (χ^2^ = 48.2, df = 1, P < 0.001).

Out of the 53 bats that were negative on J12, 32 bats (60.4%) became positive after flies were reintroduced (χ^2^ = 43, df = 1, P < 0.001). The effect of flies on prevalence was much more pronounced for bats that were born into the colony in 2010 and 2011 than for adult bats: 16/17 (94.1%) captive-born cohort bats became positive after reintroduction versus 16/36 (44.4%) wild-caught cohort bats (χ^2^ = 9.9, df = 1, P < 0.001). Including bats that were already positive on J12, 48/84 (57.1%) either became positive or changed *Bartonella* species after fly reintroduction (χ^2^ = 64.4, df = 1, P < 0.001). This effect was greater for captive-born cohort bats than for wild-caught cohort bats: 22/28 (78.6%) late cohort bats versus 26/30 (46.4%) early cohort bats (χ^2^ = 6.6, df = 1, P = 0.005). However, when comparing bats that received flies versus those that did not (i.e., cases versus controls), there were no significant differences between groups in their change in infection status after fly reintroduction (see Appendix 2 for details). Thus, the effect of bat fly reintroduction was only observable at the population-level infection prevalence and within age classes, but not for individual bats.

Bat fly reintroduction had similar effects on measures of infection load in the colony. Infection load in each sample as measured by RT-PCR cycle threshold (Ct) values (Figure 1B) and the number of positive genetic markers per sample (Figure S9A) reached a peak on M10, then declined before sharply increasing after the reintroduction of flies. This trend is reflected in the segmented regression of both measures, with a shift from positive to negative slope near day 221 and a shift from negative to positive slope near day 903 (Figure S6B,C; Table S4). Coinfection prevalence also showed a peak near M10 and declined until July 2011 (day 715) when it began to increase again (Figure S9B). However, neither of the shifts in slope for coinfection prevalence were statistically significant (Figure S6D; Table S4). For details on prevalence and load in bat flies and wild bats collected on M10 and J12, see Appendix 2.

### *Patterns of* Bartonella *diversity*

Similar to infection prevalence and load, *Bartonella* diversity measures changed in response to bat fly population density. *Bartonella* diversity was measured at two scales, at the colony level and at the individual host level. *Bartonella* species richness and evenness (Shannon index) measured colony-level alpha diversity. The number of *Bartonella* species in an individual sample and beta diversity (binomial index) measured individual-level diversity. Diversity measures showed qualitatively similar patterns during the early phase of the experiment (Figure 2A; Figure S10): an initial increase with the entry of the first three cohorts into the colony reaching a maximum in January 2010 followed by a decline. Diversity measures increased again until the reintroduction of flies on J12 and then declined slightly (or remained flat in the case of species richness). The observed trends were only partially reflected by segmented regression breakpoints. Segmented regression detected only one breakpoint each in the timelines for species richness, species evenness, and the number of *Bartonella* species in an individual sample (Table S4). A shift from positive to negative slope was detected in January 2010 for species richness (Figure S11A) whereas a change from negative to positive slope was detected for species evenness and the number of species in an individual sample between November 2010 and March 2011 (Figure S11B,C; Figure S12A). There were two significant breakpoints detected in the timeline of beta diversity, changing from negative to positive slope in July 2010 and from positive to negative slope in January 2012 (Figure S12B; Table S4). For details on diversity measures in bat flies and wild bats collected on M10 and J12, see Appendix 2.

**Figure 2.**
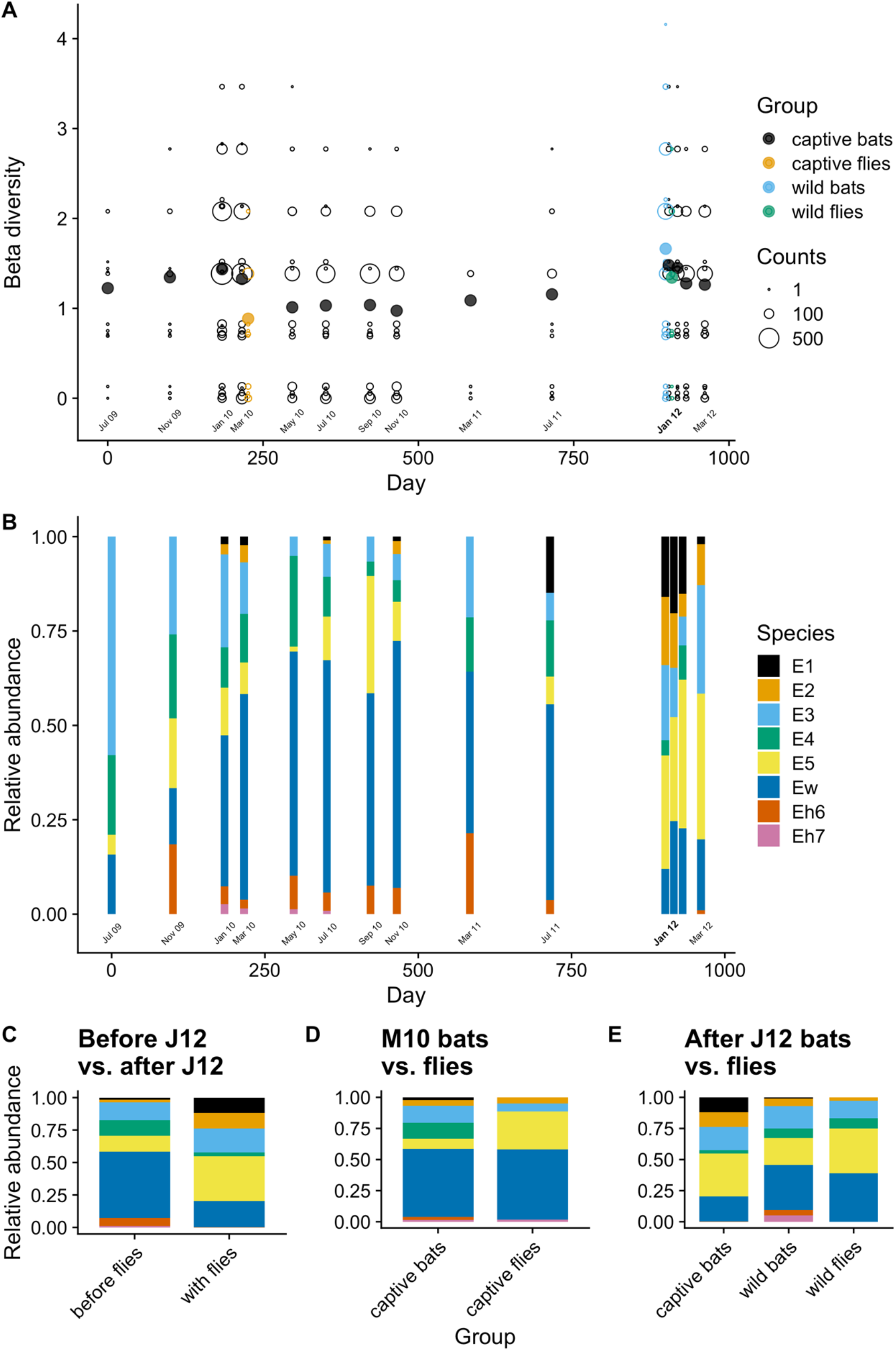
Changes in *Bartonella* beta diversity and the relative abundance of *Bartonella* species in bats and bat flies over time. Beta diversity (A) was calculated using the binomial index comparing across all infected bats and bat flies in the colony. Data for individuals are shown as open circles for each individual with the width proportional to the number of individuals with the same index value. Solid circles show the mean values. Relative abundance (B) at each time point was estimated from the total number of counts for each *Bartonella* species based on sequences from ITS, *gltA*, and *ftsZ*. For panels A and B, the month labeled in bold font on the x-axis shows when bat flies were reintroduced. Tests for differences in the relative abundance of species were performed between bats in the captive colony before and after bat flies were reintroduced on 17 January 2012 (C); between bat flies sampled from the colony and the captive bat population in March 2010 (D); and between bat flies and wild bats sampled on 17 January 2012 and the captive colony population after flies were reintroduced (E).

### *Shift in* Bartonella *species abundance*

*Bartonella* species observed in the colony varied in their relative abundance, with an apparent shift in the dominant species during the study (Figure 2B). While rarer species E1, E2, and Eh7 were not observed at all time points, E1 and E2 were consistently observed over the duration of the study. In contrast, the rarest species Eh7 was not observed after July 2010, even after flies were reintroduced to the colony. Species Eh6 was also uncommonly observed during the study, went unobserved for three time points in 2012, but was observed again in March 2012.

As noted above, beta diversity decreased after January 2010 when the bat fly population was decreasing, reached another maximum in January 2012, and then decreased again after flies were reintroduced (Figure 2A). These decreases in beta diversity correspond with periods of expansion by some species within the colony that appear to homogenize beta diversity. During a period from January 2010 to July 2011, Ew became the most abundant species in the colony (Figure 2B). Another measure of this species’ dominance in the colony is the duration of its infections in individual bats. For each individual bat that was sampled more than once and was recorded as having the same *Bartonella* species for a sequential period, we tabulated which species was present for the most time points (Figure 3). Among *Bartonella* species, Ew was the longest lasting infection in the highest number of bats (n = 40). The infection durations for this species ranged from 37 to 610 days with a median of 145 days.

**Figure 3.**
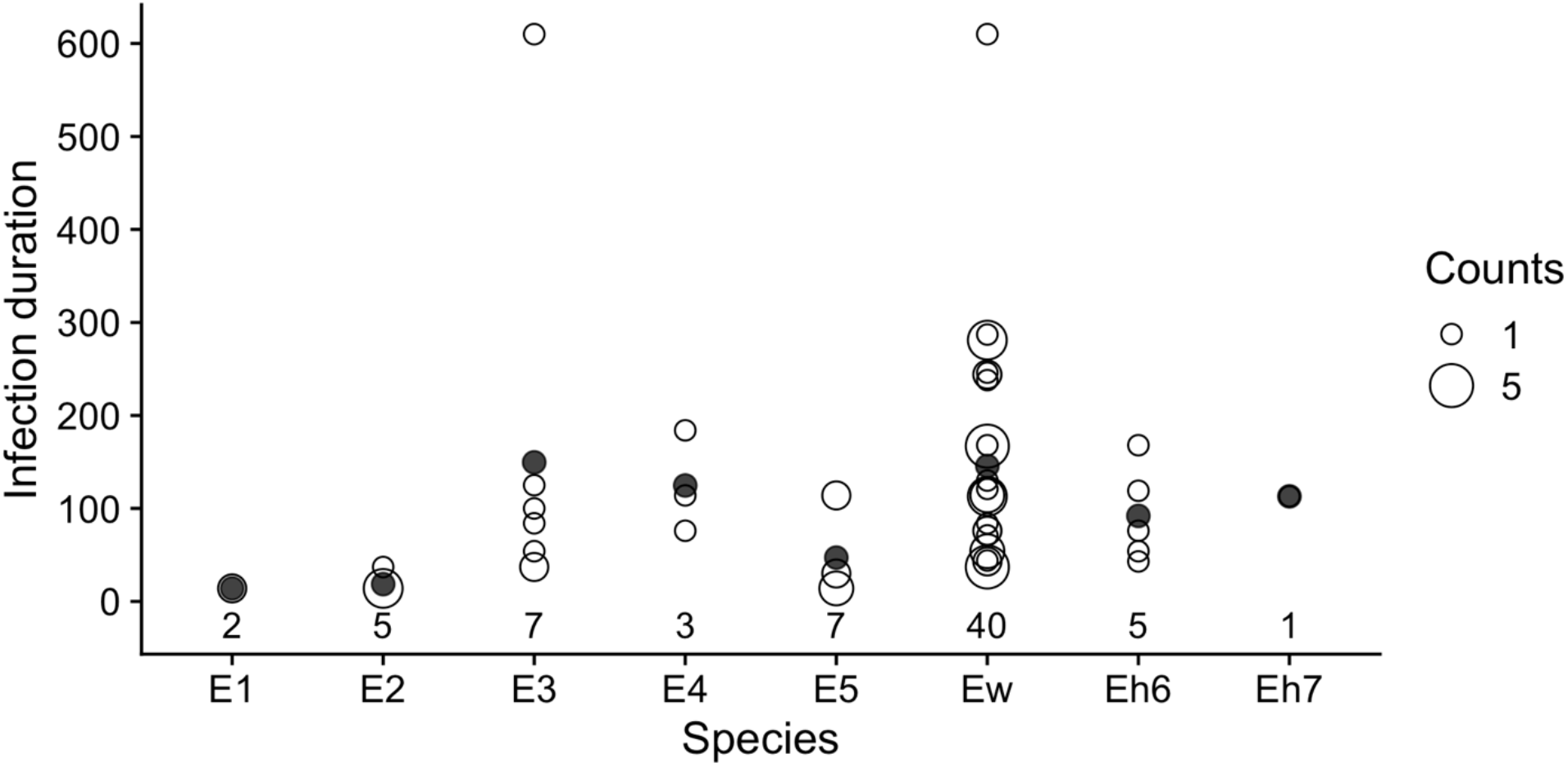
Duration of *Bartonella* sp. infections in serially infected individuals. For each *Bartonella* species, the numbers below the points are counts of individual bats that had the *Bartonella* species as its longest lasting infection (i.e., the *Bartonella* species was present for the most sequential time points). The infection durations in days for all serially infected bats are plotted as open circles with the width proportional to the number of individuals with the same infection duration. Solid circles indicate the mean duration.

Beginning around March 2011, the relative abundance of Ew began declining while species E1, E2, and E5 increased (Figure 2B). Dividing the study into two parts – before flies were introduced (July 2009 to July 2011) and after flies were introduced (J12 and after) – a clear difference in the rank abundance of *Bartonella* species was observed (Figure 2C). This shift in abundance after the introduction of flies was significant according to a multinomial LR test (D = 350.1, df = 7, P < 0.001) and individual binomial LR tests for all species (Table S5). Significant differences were also observed in the relative abundances between bat flies and sampled bat populations on M10 and J12 (Figure 2D,E; Table S6). Patterns in the occurrence of species over time and relevant tests of differences in the *Bartonella* community were similar if the relative counts (presence/absence of species across any marker rather than counts across markers) were used instead of relative abundance (Figure S13). For details on this and tests of differences in the relative abundance of species in bat flies and wild bats, see Appendix 2.

### *Interactions between* Bartonella *species*

Using multinomial and binomial LR tests on coinfection frequencies, there was evidence of both negative and positive interactions between *Bartonella* species over the period of the experiment (Figure 4). Bats infected with Ew were significantly less likely to be coinfected with E2, E3, and E5; a reciprocal negative effect on Ew from these species was not detected. Related to this, the proportion of Ew infections that were also coinfections was low (30%) considering its high relative abundance in the population over time (Figure 2B). Species E1 and Eh6 had a reciprocal negative effect on each other. Reciprocal positive effects (i.e., more coinfections than expected) were found between species E3 and E5 and species E1 and E5. Also, bats were more likely to be coinfected with Ew if they were already infected with E1, but there was no significant reciprocal effect of Ew on E1 (Figure 4).

**Figure 4.**
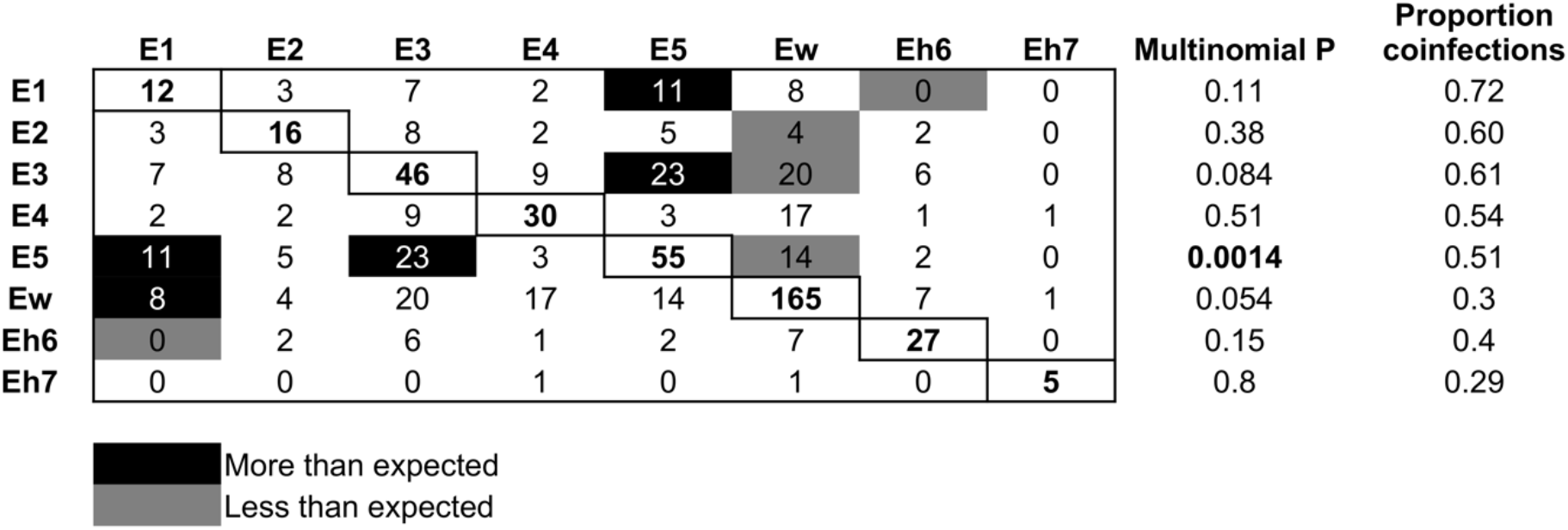
Patterns of *Bartonella* species coinfection. Rows are the focal species and columns are the partner infections. Numbers in the boxes are counts of coinfections between each pair of species; single infection counts for each species are on the diagonal. Black boxes show coinfections that occurred more frequently than expected, grey boxes show those that occurred less frequently than expected, and white boxes showed no significant pattern. Expected counts were based on the frequency of single and double infections of each *Bartonella* species, and significance was based on multinomial and binomial tests. The proportion of infections by each *Bartonella* species that were also coinfections are shown in the last column.

## Discussion

Parasites do not infect hosts in isolation but instead form diverse communities in hosts that vary over time. However, it is unclear if the same forces that affect diversity in communities of free-living organisms act in the same way or with different strengths in parasite communities. This study tested how well predictions of community ecology theory apply to host-vector-parasite systems through a unique approach that manipulated parasite dispersal among hosts within the population by changing the population density of the putative vector. Restriction of parasite dispersal minimized the effect of among-host transmission on *Bartonella* communities within individual hosts, thereby allowing the effects of ecological drift and selection on parasite community diversity to be measured. At the same time, observed trends in the prevalence and diversity of *Bartonella* infections within the colony over the course of vector population decline and reintroduction indicate that bat flies are biological vectors of *Bartonella* in bats. Overall, the experiment shows that *Bartonella* communities are affected by dispersal, drift, and selection in similar ways to free-living organisms, although numerous forms of ecological selection might be acting simultaneously.

We first hypothesized that *Bartonella* communities in the colony would respond to changes in among-host dispersal/transmission by bat flies. Specifically, we predicted that infection prevalence and diversity would decline concurrently with the bat fly population and then increase upon reintroduction of flies. The results indicate that *Bartonella* prevalence and infection load declined along with the bat fly population, then increased when flies were reintroduced in January 2012 (Figure 1). This effect was seen across the whole population but had a stronger effect on young bats born in the colony, likely attributable to their lack of prior exposure to *Bartonella* while flies were in low density. Only a few vectors of *Bartonella* bacteria have been confirmed through controlled exposure of hosts to infected vectors (Morick et al., 2013; Tsai et al., 2011). A previous study by Jardine et al. (2006) demonstrated declines in *Bartonella* prevalence after an experimental insecticide treatment reduced flea densities on Richardson’s ground squirrels (*Spermophilus richardsonii*), indicating that fleas are important vectors of *Bartonella*. Similar to this study, our results confirm that bat flies are likely vectors of *Bartonella* bacteria in bats.

*Bartonella* diversity also decreased over the corresponding period when flies were declining (Figure 2A; Figure S10). This decline may be attributed to the stochastic loss of rare species and the increase in abundance of some species, specifically Ew, through persistent infection (Figure 2B). Interestingly, all diversity measures actually increased prior to the reintroduction of flies, reaching a local peak in diversity in January 2012 before declining. This second decline could be attributed to the decline of the dominant Ew, allowing potentially latent infections by other species (E1, E2, E3, E5) to emerge as the dominant species infecting the bat population. The dominance of these species continued after flies were reintroduced and among-host transmission was restored, thus causing a short decline in diversity measures. These patterns indicate that dispersal of infections by flies is key to the long-term maintenance of *Bartonella* community diversity in bats.

While the experiment was originally designed to split bats into treatment versus control groups to assess the effect of bat fly reintroduction on changes in *Bartonella* infection status, this was not successful. Bats that received flies were not more likely to become infected or change *Bartonella* species after reintroduction. This probably occurred because bat flies did not remain on the bat they were placed on and instead moved among individuals in the colony. This would produce the poor correlation between infection status of bats and flies, as seen in the results presented and those of Becker et al. in vampire bats (2018). Nevertheless, this study establishes that the loss and reintroduction of bat fly vectors is associated with changes in *Bartonella* infection and diversity at the host population level.

We also hypothesized that limitation of parasite dispersal would result in stochastic losses of rare *Bartonella* species and changes in community beta diversity via ecological drift and shifts in the rank abundance of *Bartonella* communities due to local selection. The rarest species in the community, *Bartonella* species Eh7, was lost during the course of the study and was not restored when flies were reintroduced. This failure was likely due to a sampling effect, wherein flies carry only a subset of *Bartonella* species (Figure 2D,E), therefore limiting opportunities for effective dispersal of rare species. As noted above, beta diversity did not exhibit the expected increase when the fly population declined. Instead there was a decrease in beta diversity due to the dominance of species Ew (Figure 2A). This dominance of Ew was the most conspicuous trend in the dynamics of the *Bartonella* community over most of the study, except for the end of the experiment when there was a shift towards the next most abundant species, E5, and other lower ranked species (Figure 2B). This shift towards E5 and the decline in Ew occurred before the reintroduction of flies and was independent of the effects of among-host dispersal (due to the low density of flies at this time). We speculate that this is an emergent pattern due to within-host selection against Ew by the host immune system. Specifically, as Ew came to dominate within the population and in individual bats, it may have become the primary target of host immune responses. As Ew was eliminated, this allowed for the emergence of other latent infections within coinfected bats. Thus, without dispersal of *Bartonella* species by bat fly vectors, we speculate that ecological drift and selection by the host immune system may cause observable changes in bacterial communities.

Finally, we expected that potential interactions among *Bartonella* species would be detectable based on coinfection frequencies, providing evidence of competition or facilitation in pathogen communities. While most interactions were not significant, species Ew has negative effects on several species and typically has few coinfections (Figure 4). In contrast, positive effects were observed between species E1, E3, and E5, which show a much higher frequency of coinfection. These results indicate that parasitic bacteria like *Bartonella* do have measurable ecological interactions which are not uniformly competitive. These positive interactions could have played a role in the replacement of Ew with E5 and other species late in the study.

From just a single experiment, we can make several inferences about the ecology of *Bartonella* infections in bats. First, they can be persistent, lasting potentially hundreds of days. Other studies have alluded to the possibility of persistent *Bartonella* infection with periodic recrudesence in rodents (Bai et al., 2011; Goodrich et al., 2020; Kosoy et al., 2004) and bats (Becker et al., 2018); however, these studies were conducted in open populations where reinfection by vectors was likely frequent. Although we cannot rule out that some reinfection occurred due to the remnant bat fly population in the colony, the decline in bat fly density should have reduced reinfections relative to studies of wild populations. Second, *Bartonella* community diversity can be driven by dispersal, drift, and selection. The current study has shown that when dispersal is limited, the effects of ecological drift and selection can be more apparent. Two types of ecological selection can occur in these parasite populations, either through interactions with the host immune system or through interspecific interactions. As noted above, the immune system may lead to periodic selection against the dominant infection, a negative frequency-dependent mechanism that might help maintain diverse parasite communities (Fallon et al., 2004).

Dominance appears to be a similar facet of the composition of bacterial communities as it is in free-living organisms (Smith & Knapp, 2003). The dominance of Ew may thus stem from multiple facets of its ecology. First, it appears to be persistent within bats (Figure 3) and second, it appears to be readily taken up by flies (Figure 2D,E). We note that Ew is also the most clonal, i.e., genetically homogenous, species in the community and might be a more recently evolved or introduced species in *E. helvum* (Bai et al., 2015). While there was no evidence that Ew caused higher infection loads (by Ct value or number of markers positive), the resolution of our sampling protocol probably was not high enough to detect this. Future studies should inspect growth curves throughout the infection cycle to see if Ew has any growth advantage. Other forms of interference or resource competition must be explored further, perhaps through controlled infection experiments.

Future work within this system might involve controlled exposure of *Bartonella*-negative bats and confirmation of the exposure route. Alternative routes might include bat fly bite, requiring tropism of the bacteria to the salivary glands, or contamination through bat fly feces, requiring replication in the fly gut and persistent shedding of viable bacteria in feces. Additional studies could examine immune function in bats (Boughton et al., 2011) in response to *Bartonella* infection to confirm the existence of frequency-dependent selection against *Bartonella* species and to help determine the appropriate epidemiological models to explain *Bartonella* infection dynamics (Brook et al., 2017).

This study has contributed to a more comprehensive understanding of the ecology of *Bartonella* species in bats and connects with broader community ecology theory developed in free-living and symbiotic organisms (Costello et al., 2012; Miller et al., 2018; Vellend, 2010). Limitation of dispersal in this experiment led to declines in local species diversity in individual bats, a pattern that fits well with predictions from patch dynamics or mass effects models of metacommunities (Leibold et al., 2004). The results also show that not all bacterial interactions are negative, even those that presumably share the same niche. This parallels the recognized importance of positive species interactions in plant communities (Bertness & Callaway, 1994) and among bacterial taxa in animal microbiomes and aquatic habitats (Faust et al., 2012; Hegde et al., 2018; Ju & Zhang, 2015). A recent study by Gutiérrez et al. (2018) on *Bartonella* infections in desert rodents showed a mixture of negative, neutral, and positive interactions similar to the present study. Theoretical and experimental studies suggest that communities remain stable through a predominance of neutral or weak species interactions that can attenuate large competitive or facultative effects (Aschehoug & Callaway, 2015; McCann, 2000). Weak interactions, paired with the frequency-dependent selection discussed above, could provide a model for understanding how *Bartonella* species and other parasitic microorganisms coexist in communities within their hosts. Such mechanisms could allow bacteria to share a niche or split it temporally, which could lead to periodic shifts in the dominant species but maintain the community as a whole. Future work using this system and similar longitudinal studies on other pathogens in natural host populations could lead to additional insights on the nature of microorganismal communities and the broad ecological processes that act across taxonomic and spatial scales.

## Acknowledgements

We acknowledge the help of many people who assisted with maintenance of the bat colony and sampling during this study, including staff of the Accra Zoo (Wildlife Division of the Forestry Commission of Ghana) and other researchers who assisted with the work. We also thank current and former members of the Webb Lab for their comments on early versions of the manuscript. AAC was supported by a Royal Society Wolfson Research Merit award. JLNW is supported by the Alborada Trust. DTSH received funding from the Wellcome Trust and the Royal Society Te Apārangi (MAU1701).

## Data accessibility

The data that supports the findings of this study are available in the supplementary material of this article. Representative sequences for *Bartonella* genogroups Eh6 and Eh7 and two *Rickettsia* sequences from *E. helvum* and *C. greefi* have been submitted to GenBank under the accession numbers MN249715–MN249720, MN250730–MN250788, and MN255799–MN255800. Phylogenetic trees, R code, and additional data sheets will be made available on Dryad.

## Author contributions

DTSH, JLNW, AAC, YN, and RS designed research; CDM, MYK, YB, LMO, RS, and DTSH performed research; CDM analyzed data; CDM, CTW, and DTSH wrote the paper; all authors contributed to and approved the final version of the paper.

