## Supplemental Information for "Manipulating vector transmission reveals local processes in bacterial communities of bats"

**Supplemental Information for:**
**Manipulating vector transmission reveals local processes in bacterial communities of bats**

Clifton D. McKee, Colleen T. Webb, Michael Y. Kosoy, Ying Bai, Lynn M. Osikowicz, Richard Suu-Ire, Yaa Ntiamoa-Baidu, Andrew A. Cunningham, James L. N. Wood, David T. S. Hayman

### Appendix 1. Supplementary methods

*Sampling protocol*

Captive bats (Table S1) were collected using hand nets after cordoning bats into one quarter of the cage with a curtain system and then placed into a smaller cage until processing. Wild bats (the first three captive cohorts and those from 31 January 2012) were captured from roosts using 6–18 m mist nets or hand nets then placed in individual cloth bags until processing. While under manual restraint, the surface of the inner wing along the propatagial vein was wiped with 70% ethanol then 0.2–1.0 ml of whole blood was collected using a citrated 1 ml syringe and transferred into labeled microcentrifuge tubes. After bleeding had ceased, bats were released either into the main area of the enclosure or back to the wild roost. On two occasions bat flies were collected from bats, M10 from the captive colony and J12 from the wild source colony. Bat flies were collected from the pelage of bats and placed in individual sterile tubes labeled with the ID number of the host bat. Whole blood was immediately frozen at -80 ºC or blood clots were separated from serum then frozen at -80 ºC; bat flies were kept frozen at -80 ºC. All samples were shipped to the Centers for Disease Control and Prevention Division of Vector-Borne Diseases in Fort Collins, CO on dry ice where they were kept at -20 ºC or below until DNA extraction.

*DNA extraction*

Bat flies were rinsed in 70% ethanol then in sterile 1x PBS (0.15 M, pH 7.5, CDC, Atlanta, GA) before being transferred to a clean microcentrifuge tube and triturated in 500 µl brain heart infusion (BHI) broth (CDC, Atlanta, GA) using a sterile pestle. Genomic DNA was extracted from bat blood samples and triturated bat flies either by hand using a QIAamp DNA Mini Kit (Qiagen, Valencia, CA) or with a QIAxtractor automated instrument (Qiagen) following the manufacturer's protocols for tissues (flies) and blood. Extraction controls (blank wells) were included to ensure no cross-contamination occurred during extraction. Extracted DNA was stored in clean microcentrifuge tubes at -20 or 4 °C during the duration of pathogen testing.

*Bacterial detection and sequencing*

*Bartonella* spp. were detected via conventional PCR targeting the 16S–23S intergenic spacer region (ITS) via single-step PCR (Diniz et al., 2007) and the citrate synthase (*gltA*) and cell division protein (*ftsZ*) genes via nested PCR (Bai et al., 2016). Quantification of *Bartonella* infection load was performed using real-time PCR targeting the *Bartonella* transfer-messenger RNA gene (*ssrA*) (Diaz et al., 2012). *Rickettsia* DNA was detected using a separate real-time PCR assay targeting the 23S ribosomal RNA subunit (Kato et al., 2013). Samples positive for *Rickettsia* by real-time PCR were confirmed using conventional PCR targeting the *Rickettsia gltA* gene using a nested protocol (Choi et al., 2005; Lee et al., 2014; Regnery et al., 1991). Primers and thermocycler protocols for all real-time and conventional PCR are listed in Tables S2–S3.

All PCR amplifications were run in a C1000 Touch Thermal Cycler (Bio-Rad, Hercules, CA) with the addition of a CFX96 Real-Time System (Bio-Rad) for real-time PCRs. For all

PCRs, positive (*Bartonella doshiae*, *Rickettsia felis*) and negative (RNase-free water only) controls were included to determine correctly sized amplicons and to detect potential cross-contamination, respectively. PCR products were inspected for the presence of positive amplicons of the correct size by gel electrophoresis using 1.5% agar and GelGreen stain (Biotium, Hayward, CA). Positive amplicons were purified using a QIAquick PCR Purification Kit (Qiagen) and sequenced in both directions using an ABI 3130 Genetic Analyzer (Applied Biosystems, Foster City, CA). Forward and reverse reads were assembled and edited using the SeqMan Pro program in Lasergene v14 (DNASTAR, Madison, WI). Sequences obtained from bat blood or bat flies were initially confirmed to the bacterial genus using the Basic Local Alignment Search Tool (BLAST; https://blast.ncbi.nlm.nih.gov/Blast.cgi).

*Phylogenetic analysis of bacterial sequences*

Bacterial sequences obtained from bats and bat flies were aligned with reference sequences for named species from each detected bacterial genus and with sequences for each bacterial genus that have been detected previously in bats (Supplementary Data). Alignments for each genetic locus were performed separately using MAFFT v7.471 (Katoh & Standley, 2013) using the L-INS-i method. Alignments were trimmed to a common length eliminating poorly aligned positions using Gblocks v0.91b (Castresana, 2000). Alignments were then visually inspected for errors and manually corrected. Concatenation of multiple loci for phylogenetic analysis was performed after alignment and trimming using Phyutility v2.7.1 (Smith & Dunn, 2008). Simultaneous model selection and maximum likelihood reconstruction of phylogenetic trees was performed using IQ-TREE v2.1.1 (Kalyaanamoorthy et al., 2017; Minh et al., 2020; Nguyen et al., 2015). Branch support for each tree was estimated from 1000 ultrafast bootstrap samples from the respective alignment (Hoang et al., 2018).

*Regression analyses*

Linear regression was performed to determine demographic factors that influence *Bartonella* infection status for bats sampled from the colony in March 2010. Two data sets were used: a set containing data from all bats and a set containing data from females only. In the full data set, covariates included bat sex, age class (neonate, juvenile, sexually immature adult, and sexually mature adult) following Peel et al. (2016) and the presence/absence of bat flies on each bat. In the females only data set, covariates were the same as the full dataset (excepting sex) but also included pregnancy status (pregnant or not). Data were fit to covariates in a generalized linear model (GLM), treating infection status as a binomial variable with a logit link. Model selection was then performed based on Akaike's information criterion corrected for finite sample sizes (AICc) using the dredge function in the R package *MuMIn* (Bartoń, 2020; R Core Team, 2020). The model with the smallest AICc was chosen unless another model was less than two AICc points away from the top model (Burnham & Anderson, 2004), in which case the simpler model was chosen.

Segmented regression was performed to detect breakpoints in measures of *Bartonella* prevalence and diversity over the course of the experiment. Separate GLMs were fit for each measure in R. *Bartonella* single infection and coinfection prevalence were both treated as binomial variables with a logit link. Real-time PCR cycle threshold (Ct) values, Shannon number, and inverse Simpson index were treated as gamma-distributed variables. The number of sequenced *Bartonella* markers, *Bartonella* species richness, and number of *Bartonella* species in a sample were treated as Poisson-distributed variables. Segmented regression was performed on fitted GLMs using the R package *segmented* (Muggeo, 2020) with breakpoints estimated with 1000 bootstrap iterations.

*Likelihood ratio tests*

We tested whether *Bartonella* species coinfected bats with other species more than expected by chance using a multinomial test adapted from a previous study analyzing patterns of influenza A transmission in birds (Pepin et al., 2013). Following Pepin et al., only double infections (two coinfecting species) were included because higher order infections were rare and challenging to interpret. The null hypothesis for the test was that *Bartonella* species $i$ would coinfect with any other species $j$ with equal probability and the expected counts for partner coinfections of species $i$ would be proportional to the frequency of each partner in all single (*s*) and double (*d*) infections. Therefore, the expected counts for each $j$ partner of species $i$, $E[X_j]$ are: $E[X_j] = (X_j^{s+d}/N_j^{s+d}) * X_i^d$, where $X_j^{s+d}$ is the total number of single and double infections for partner $j$, $N_j^{s+d}$ is the total number of single and double infections for all $j$ partners, and $X_i^d$ is the total number of double infections for species $i$. The maximum likelihood estimates for the parameters in the null multinomial model for each species $i$ are then: $\pi_j = E[X_1]/\sum(E[X_j]), \ldots, E[X_N]/\sum(E[X_J])$. The probabilities under the null and alternative models are: $P(X)_0 = N_j^d!\prod(\pi_j^{xi}/X_i!)$ and $P(X)_A = N_j^d!\prod(p_j^{xi}/X_i!)$ and the likelihood ratio statistic $D$ is $-\ln(P(X)_0/P(X)_A)$, which is approximately distributed $\chi^2_{n-1}$. The likelihood ratio statistic was divided by the correction factor $1 + \sum(\pi_j^{-1} - 1)/6N_j^d(n-1)$ to decrease type I error inflation due to the difference between the moments of the likelihood ratio statistic and the chi-square distribution. Differences between the observed and expected counts of coinfections were tested using binomial likelihood ratio tests, using the same correction factor as above. Functions for multinomial and binomial likelihood ratio tests were written in R. These functions were first used to test for differences in observed and expected counts of coinfections for the whole course of the experiment (961 days). Additional tests were performed on two partitions of the experiment: bats sampled before J12 and bats sampled after J12. This was based on visual observation of a change in the frequency of *Bartonella* species starting around this point in the experiment.

In addition to the tests of the observed and expected counts of coinfections, these same likelihood ratio test functions were used to perform tests on changes in the frequency of single infections and coinfections in the captive colony over time and differences in the relative frequency of infections between bats and bat flies. Specifically, we performed likelihood ratio tests on the relative frequency of *Bartonella* species before versus after J12, using the before-J12 frequencies as the expected frequencies to calculate the likelihood ratio statistic. We calculated the differences in the frequency of *Bartonella* species in bats versus bat flies sampled on M10 and J12, using the frequencies in bats as the expected frequencies. We also calculated the differences in the frequencies of *Bartonella* species in bats after J12 versus bat flies sampled on J12, again using the frequencies in bats as the expected frequencies.

*Assumptions*

Within this system, *Bartonella* infection does not cause obvious signs of disease in bats or flies (Kosoy et al., 2010), so we assume that there are no parasite-mediated mortality effects. We consider hosts as discrete patches containing parasite species and the dynamics of these infections are linked through transmission by bat flies as they disperse among hosts. We consider *Bartonella* species as static and not measurably evolving over the current study, an assumption supported by the very low mutation rates (Gutiérrez et al., 2018). Finally, although vertical transmission of *Bartonella* from dam to offspring is possible (Kosoy et al., 1998), it has not been demonstrated in bats, so we assume that bats are born uninfected, and the primary transmission route is through vector transmission.

**Appendix 2. Supplementary results**

*Phylogenetic analysis of detected* Bartonella *and* Rickettsia *species*

*Bartonella* sequences from *E. helvum* and *C. greefi* predominantly grouped closely with six *Bartonella* species previously described from *E. helvum*: *Bartonella* spp. E1–E5 and Ew (Bai et al., 2015; Kosoy et al., 2010). The phylogenetic distinctiveness of these species can be observed based on all loci sequenced: *gltA*, *ftsZ*, and ITS (Figures S1–S3). In addition to these six species, two novel *Bartonella* genogroups were observed in both *E. helvum* and *C. greefi*, denoted *Bartonella* spp. Eh6 and Eh7. *Bartonella* sp. Eh6 was detected at all three sequenced loci whereas species Eh7 was only detected at *gltA* and *ftsZ* (Figures S1–S3). Sequences representing these two potentially novel *Bartonella* species have been submitted to GenBank with the following accession numbers: MN250730–MN250774 (*gltA*), MN250775–MN250788 (*ftsZ*), and MN249715–MN249720 (ITS).

All *gltA* sequences from species Eh6 were found to be similar to each other (88.5–100% sequence identity), and according to BLAST search, similar (88.2–94.7% sequence identity) to a sequence obtained from *C. greefi* collected from *E. helvum* on Bioko island in the Gulf of Guinea (GenBank accession number JN172066) by (Billeter et al., 2012). All *ftsZ* sequences were highly similar to each other (98.5–100% sequence identity), as were ITS sequences (99.4–100% sequence identity). All *gltA* sequences from species Eh7 were similar to one another (99.7–100% sequence identity) and similar (99.7–100% sequence identity) to six sequences (GenBank accession numbers JN172046, JN172050, JN172053, JN172058, JN172067, and JN172072) from *C. greefi* collected from *E. helvum* on Annobón and Bioko islands in the Gulf of Guinea and in Ghana (Billeter et al., 2012). Sequenced loci grouped species Eh6 and Eh7 as monophyletic groups distinct from other *E. helvum*-associated *Bartonella* species with strong bootstrap support (Figures S1–S3).

A maximum likelihood tree produced from concatenated *ftsZ* and *gltA* sequences from known *Bartonella* species and *Bartonella* strains detected in bats (Supplementary Data) demonstrates that *Bartonella* species from *E. helvum* and *C. greefi* are broadly distributed in the *Bartonella* phylogeny (Figure S4). *Bartonella* sp. Ew clusters with 100% bootstrap support with three other *Bartonella* strains isolated from *Myotis blythii* and *Rhinolophus ferrumequinum* in Georgia (Urushadze et al., 2017). *Bartonella* spp. E3, E1, E2, and E5 are part of a large and distinct clade of bat-associated *Bartonella* strains isolated from hosts in several bat families including Hipposideridae, Rhinolophidae, Miniopteridae, Emballonuridae, and Vespertilionidae in Africa and Eurasia (Kosoy et al., 2010; Lilley et al., 2015; Lin et al., 2012; McKee et al., 2017; Urushadze et al., 2017). While this clade only received 75% bootstrap support in the current tree using concatenated *ftsZ* and *gltA*, a previous analysis using three additional loci and a Bayesian phylogenetic approach found 100% posterior support for this clade (McKee et al., 2017). *Bartonella* sp. Eh6 is forms a clade including strains from *Pipistrellus pipistrellus* and *Myotis blythii* from Georgia (Urushadze et al., 2017) with 97% bootstrap support. The Bayesian analysis by McKee et al. (2017) showed that this smaller clade is included as a subclade within the larger Old World bat-associated clade mentioned above with 100% posterior support. *Bartonella* spp. E4 and Eh7 were found to be most closely related to each other, although with only 51% bootstrap support. These two species are included in a larger clade containing *Bartonella* strains associated with rodents, carnivores, marsupials, and another bat (*Myotis emarginatus*) from Georgia (Urushadze et al., 2017). While the bootstrap support for this larger clade is low (58%), sequencing of additional markers may result in higher support (McKee et al., 2017; Urushadze et al., 2017).

Only one bat and one fly were positive for *Rickettsia* sp. DNA, both sampled on M10; however, the positive fly was not collected from the positive bat. The two *Rickettsia gltA* sequences obtained from *E. helvum* and *C. greefi* in March 2010 were identical to one another (313/313 bp). Both sequences have been submitted to GenBank with accession numbers MN255799 and MN255800. A maximum likelihood tree generated from *gltA* sequences showed that these *Rickettsia* sequences are distinct from those previously obtained from bats or bat ectoparasites (Figure S5). The sequences were most closely related to *Rickettsia akari* (89% bootstrap support) within the transitional group rickettsiae clade which also includes *R. felis*, *R. hoogstraalii*, *R. lusitaniae*, and *R. australis* (Sánchez-Montes et al., 2016; Weinert et al., 2009). Other bat-associated *Rickettsia* strains have been detected from this clade in mainly insectivorous bats or their associated soft ticks in Africa, Eurasia, and North America (Hornok et al., 2018, 2019; Sánchez-Montes et al., 2016).

*Demographic patterns of* Bartonella *prevalence during experiment*

We observed no significant differences in the proportion of males and females infected (Figure S8B) at the start of the study ($\chi^2 = 1.1$, df = 1, P = 0.29) or by the end of the study ($\chi^2 = 0.17$, df = 1, P = 0.68). However, there was a significant increase in the proportion infection between the start and end for males ($\chi^2 = 19$, df = 1, P < 0.001) and females ($\chi^2 = 10.5$, df = 1, P < 0.001), although this is linked to the increases observed in neonates/juveniles and adults.

*Effects of bat fly reintroduction on treatment versus control bats*

Despite the significant changes in prevalence and infection load observed after J12, the reintroduction of flies into the colony was intended to be a randomized treatment/control study to compare bats receiving flies to those that did not receive flies in terms of their change in infection status. For all bats that tested negative for *Bartonella* on J12, bats that received flies were not more likely to become infected than bats that did not receive flies ($\chi^2 = 0.012$, df = 1, P = 0.54). This pattern remains even if bats were split into two groups: sexually immature adults and adults ($\chi^2 = 0.25$, df = 1, P = 0.31) and neonates and juveniles ($\chi^2 = 2.5E{-}31$, df = 1, P = 0.5). Including the bats that were positive for *Bartonella* on J12, bats that received flies were slightly more likely to become infected or change *Bartonella* species than bats without flies, but this difference was not significant when all age groups were combined ($\chi^2 = 1.5$, df = 1, P = 0.11). However, sexually immature and sexually mature adult bats that received flies were more likely to become infected or change *Bartonella* sp. than bats that did not receive flies ($\chi^2$ = 3.2, df = 1, P = 0.036). A similar pattern was not observed in neonates and juveniles ($\chi^2$ = 0.38, df = 1, P = 0.27).

Additionally, there was poor correspondence between the *Bartonella* species found in the colony bats that received flies with the *Bartonella* species found in the bats that were the donors for the flies or other flies that were removed from the donor bats. The frequency of finding the same *Bartonella* species in the recipient bat and either the donor bat or a sampled fly taken from the donor bat (13/27, 48.1%) was no better than random ($\chi^2$ = 5E-31, df = 1, P = 0.5). This was again true if bats were subdivided into sexually immature and sexually mature adults (5/13, 38.5%; $\chi^2$ = 0.039, df = 1, P = 0.58) and neonates and juveniles (8/14, 57.1%; $\chi^2$ = 2.8E-32, df = 1, P = 0.5). Using the additional data from the collection of bat flies on M10, we also observed no correlation between the presence of a fly and whether a bat was positive (Pearson's R = -0.067, t = -0.52, df = 59, P = 0.61). The frequency of finding the same *Bartonella* species in the bat and the sampled bat fly (9/26, 34.6%) was no better than random ($\chi^2$ = 0.71, df = 1, P = 0.8).

*Differences in* Bartonella *prevalence and diversity between bats and bat flies*

*Bartonella* prevalence in bat flies (93%) collected from the colony on M10 was similarly high as in the colony bats (Figure 1A). The flies collected on J12 from the wild bat population had a slightly lower infection prevalence (89%) compared to the wild bats (94%), and both the wild flies and wild bats had higher prevalence than the bats in the colony (31%) on the same date. Average infection loads in flies on M10 were less than in the colony bats, indicated by higher Ct values (Figure 1B). Similarly, wild bat flies had higher Ct values on J12 than the wild bats but were lower than in the colony bats. Bat flies had a higher average number of positive markers but lower coinfection prevalence compared to bats from their respective populations in the colony on M10 and from the wild population on J12 (Figure S9).

On M10, all *Bartonella* diversity measures (beta diversity, species richness, Shannon index, inverse Simpson index, number of species an individual sample) in flies were lower than in the bat population at that time (Figure 2A; Figure S10). On J12, all diversity measures except inverse Simpson index in wild bats were higher than in the captive colony. Diversity measures in flies sampled from wild bats at this time were lower than in the wild population.

There were significant differences in the relative abundance of *Bartonella* species in the bats and flies sampled on M10 (D = 43.7, df = 7, P < 0.001) with significant differences observed in species E4 and E5 using binomial LR tests (Figure 2D; Table S6). Differences between the relative abundance of *Bartonella* species after the reintroduction of flies on J12 and the wild flies that were introduced into the colony were observed (D = 16.3, df = 6, P = 0.012), with substantially higher abundance of Ew and lower abundance of E1 in the flies than the colony bats (Figure 2E; Table S6). Similarly, differences were observed in distribution of species between the wild bats and wild flies sampled on J12 (D = 16.7, df = 7, P = 0.019), with a higher abundance of E5 and lower abundance of Eh6 and Eh7 in the flies than in the wild bats (Figure 2E; Table S6). As detailed below, similar results were observed if the relative counts were used instead of relative abundance.

*Shift in* Bartonella *community diversity using relative counts*

Tests for changes in *Bartonella* diversity were initially performed using the relative abundance of *Bartonella* species based on the individual number of sequences acquired for each *Bartonella* species in a sample across the three different genetic markers. Using just the presence of a *Bartonella* species in a sample by any one of the different markers, what we term relative counts, we observed very similar patterns in the change in the distribution of *Bartonella* species over time (Figure S13) and similar statistical test results for the comparison of species distributions before and after the reintroduction of flies (Table S7). In fact, there is a very strong positive correlation (Pearson's $R = 0.99$, $t = 20.9$, $df = 6$, $P < 0.001$) between the abundance and counts for each *Bartonella* species over the entire study and at each sample time point (Figure S14).

*Individual infection histories and duration of infections*

Individual infection histories for all 112 identified bats and relevant statistics for their histories are included in the Supplementary Data. Out of the 112 individual bats sampled during this study period, 102 (91.1%) were sampled at least two time points in a row. The remaining 10 bats were either euthanized after first sampling (n = 1), were found dead between sampling time points (n = 5), or had disappeared and were presumed dead (n = 4).

There was considerable individual variation among bats in their infection histories, with some bats never becoming infected, bats with intermittent infections throughout the study, bats clearing infection soon after entry into the colony, and other bats with highly persistent infections. Of the 112 bats that were sampled, 100 (89.3%) tested positive at least once during the study and 65 (58%) bats were positive at entry into the colony. Of the 102 bats that were sampled more than once, 95 (93.1%) tested positive at least once: 80/95 (84.2%) were positive at more than one time point and 15/95 (15.8%) were positive only once. Of the 15 positive only one time, 12 (80%) were bats born into the colony in April 2010 (n = 2) or April 2011 (n = 10) and 10/12 (83.3%) became positive only after the flies were reintroduced on J12. The three adult bats only infected once were from the cohort that entered the colony in January 2010. One of these bats was positive on entry and was found dead in May 2010, another became positive shortly after entry in March 2010, and the third adult did not become positive until after the fly reintroduction.

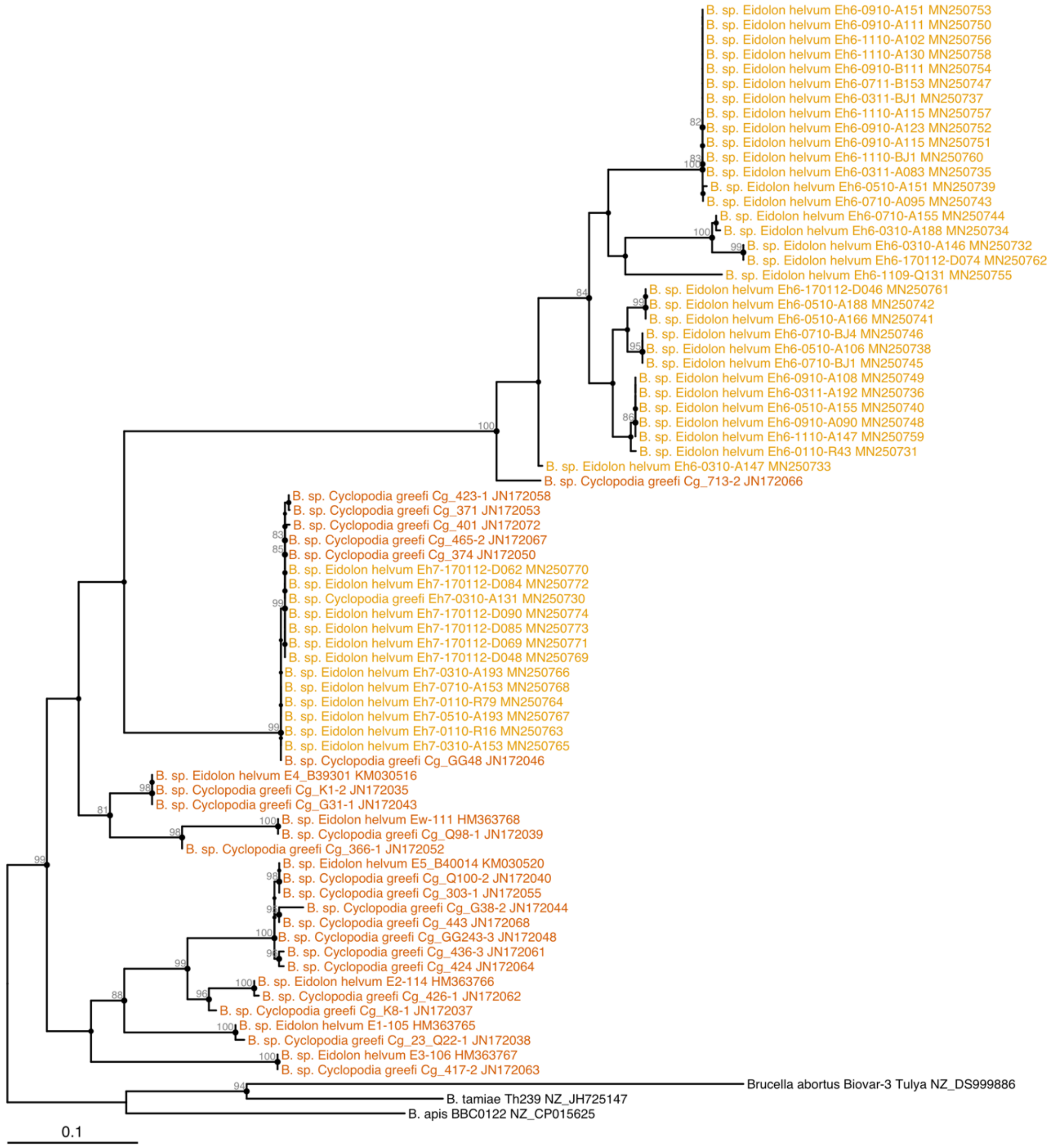


**Figure S1.** Maximum likelihood phylogenetic tree of *Bartonella gltA* sequences produced from a 356 bp alignment of 76 sequences. The best model of sequence evolution was TIM3+F+I+G4 based on AICc. The tree was rooted at the midpoint and bootstrap branch support values greater than 80% are shown in gray next to branches. Names of *Bartonella* sequences previously obtained from *E. helvum* or *C. greefi* are colored orange while names of new sequences from these species are colored yellow.

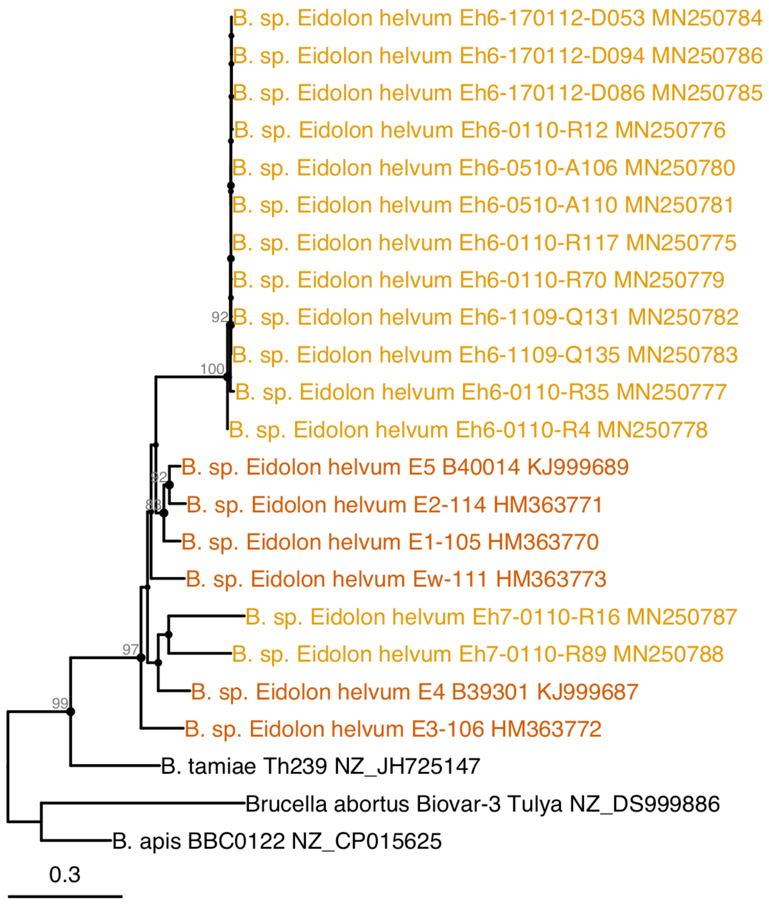


**Figure S2.** Maximum likelihood phylogenetic tree of *Bartonella ftsZ* sequences produced from a 727 bp alignment of 23 sequences. The best model of sequence evolution was TVM+F+I+G4 based on AICc. The tree was rooted at the midpoint and bootstrap branch support values greater than 80% are shown in gray next to branches. Names of *Bartonella* sequences previously obtained from *E. helvum* or *C. greefi* are colored orange while names of new sequences from these species are colored yellow.

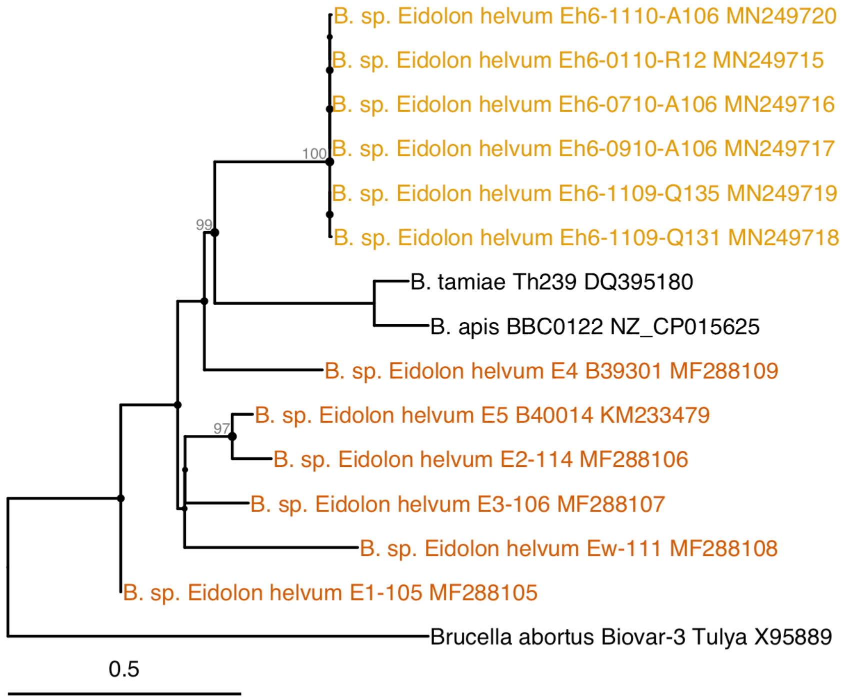


**Figure S3.** Maximum likelihood phylogenetic tree of *Bartonella* ITS sequences produced from a 446 bp alignment (including gaps) of 15 sequences. The best model of sequence evolution was TIM3+F+I+G4 based on AICc. The tree was rooted at the midpoint and bootstrap branch support values greater than 80% are shown in gray next to branches. Names of *Bartonella* sequences previously obtained from *E. helvum* or *C. greefi* are colored orange while names of new sequences from these species are colored yellow.

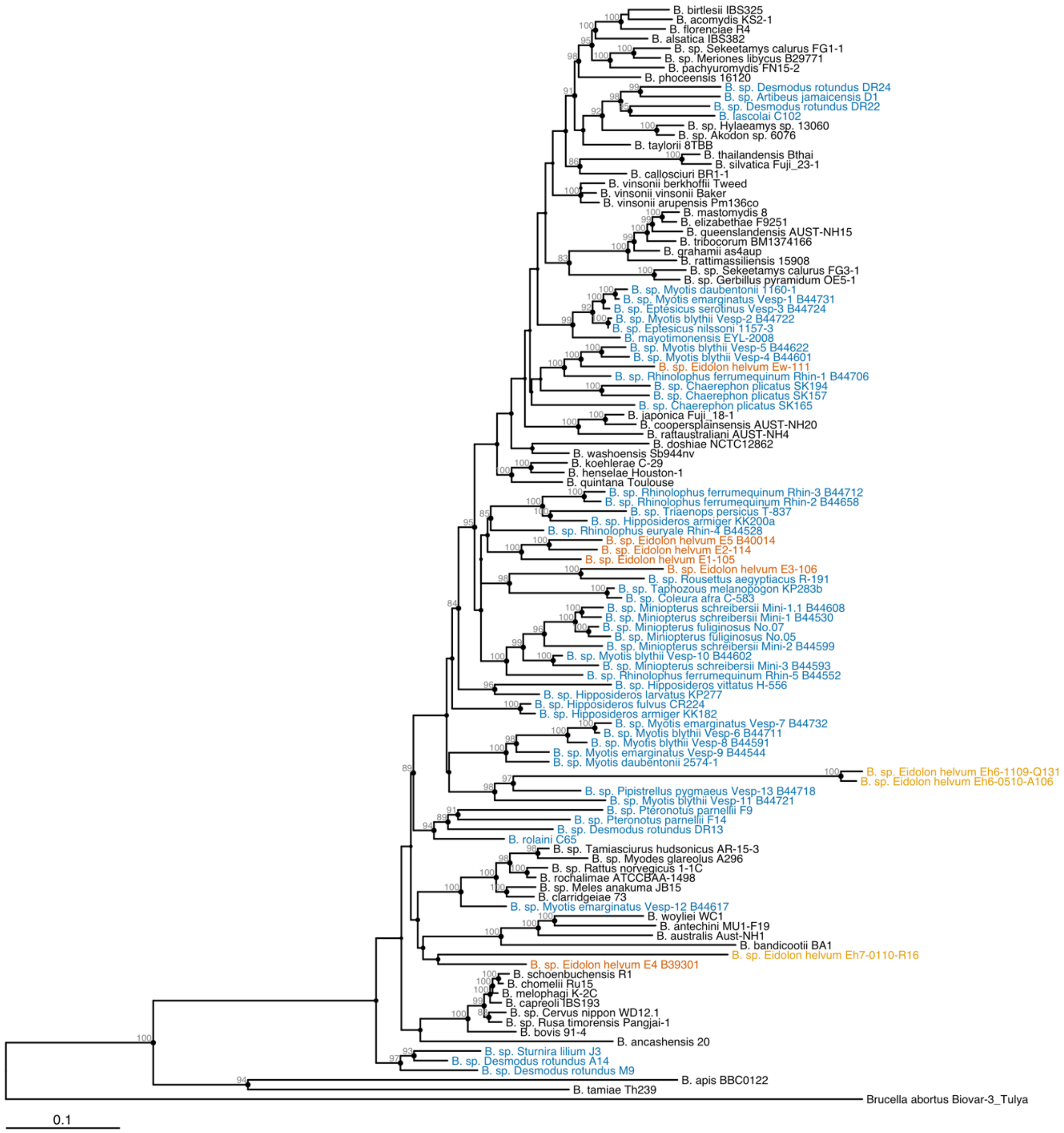


**Figure S4.** Maximum likelihood phylogenetic tree of concatenated *Bartonella ftsZ* and *gltA* sequences produced from a 1247 bp alignment (891 bp *ftsZ*, 356 bp *gltA*) of 114 sequences. The best model of sequence evolution was TVM+F+R7 based on AICc. The tree was rooted at the midpoint and bootstrap branch support values greater than 80% are shown in gray next to branches. Names of *Bartonella* species/strains previously obtained from bats are colored blue, sequences previously obtained from *E. helvum* or *C. greefi* are colored orange, and names of new sequences from these species are colored yellow.

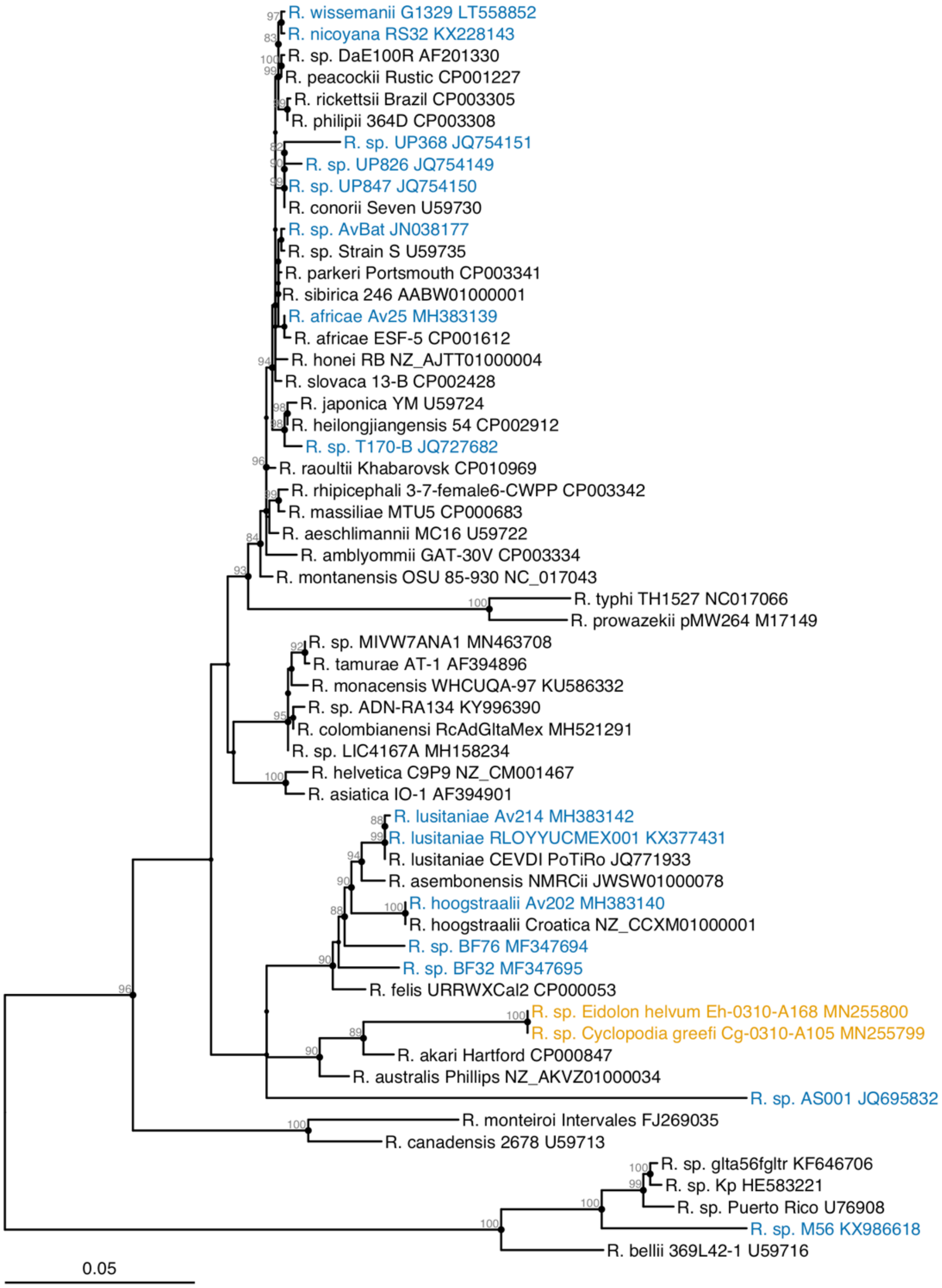


**Figure S5.** Maximum likelihood phylogenetic tree of *Rickettsia gltA* sequences produced from a 1234 bp alignment of 58 sequences. The best model of sequence evolution was TIM+F+R2 based on AICc. The tree was rooted at the midpoint and bootstrap branch support values greater than 80% are shown in gray next to branches. Names of *Rickettsia* species/strains previously obtained from bats are colored blue while names of new strains from *E. helvum* and *C. greefi* are colored yellow.

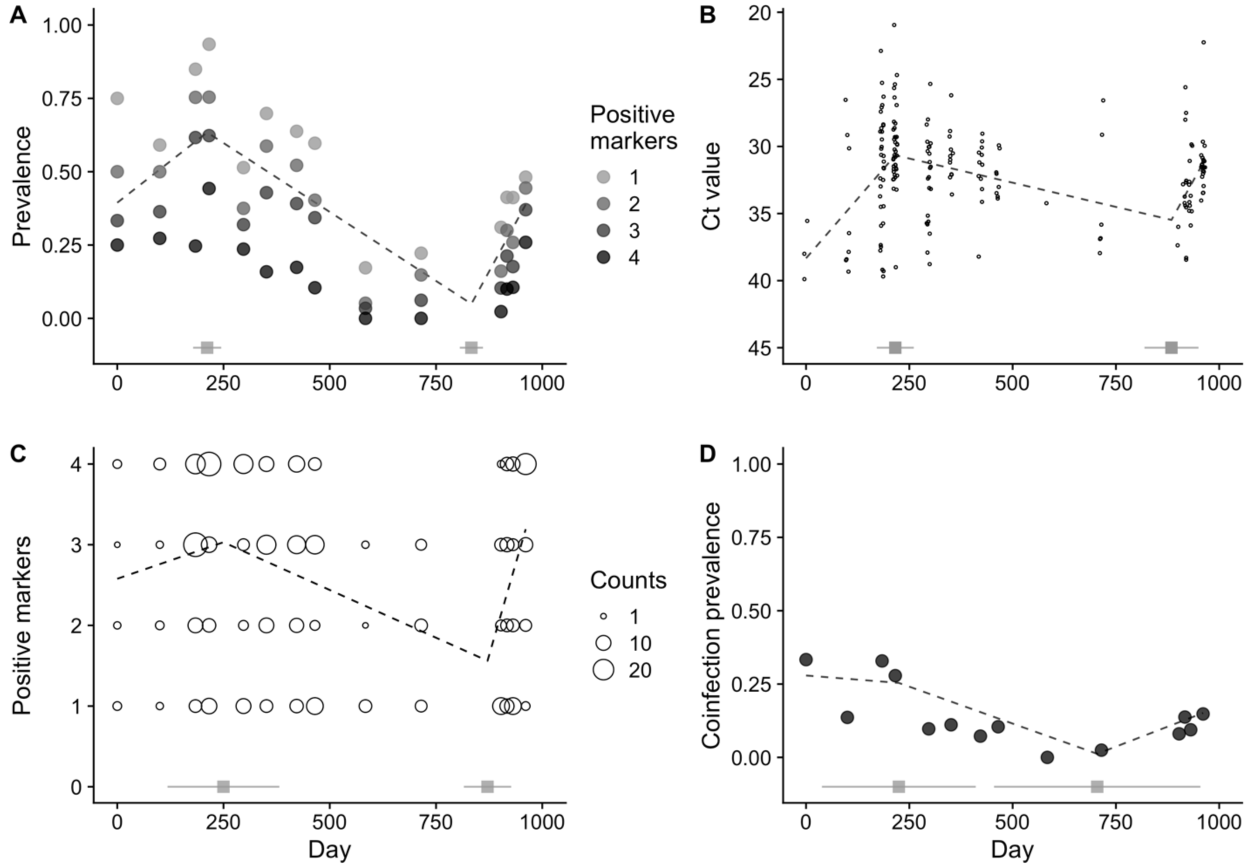


**Figure S6.** Segmented regression analysis of *Bartonella* prevalence and load. (A) Points for *Bartonella* prevalence are shown considering one or more, two or more, three or more, or all four markers positive (including RT-PCR). (B) Only points with RT-PCR Ct values < 40 are shown. (C) Points show the number of markers that were positive for each individual with the width proportional to the number of individuals positive at that many markers. (D) Coinfection prevalence was measured by the number of individuals that were positive for two or more *Bartonella* species at each time point. For each measure, dashed lines for the predicted trend from segmented regression are drawn over the data points. Breakpoints and 95% confidence intervals estimated by segmented regression are shown above the x-axis as grey squares and solid lines.

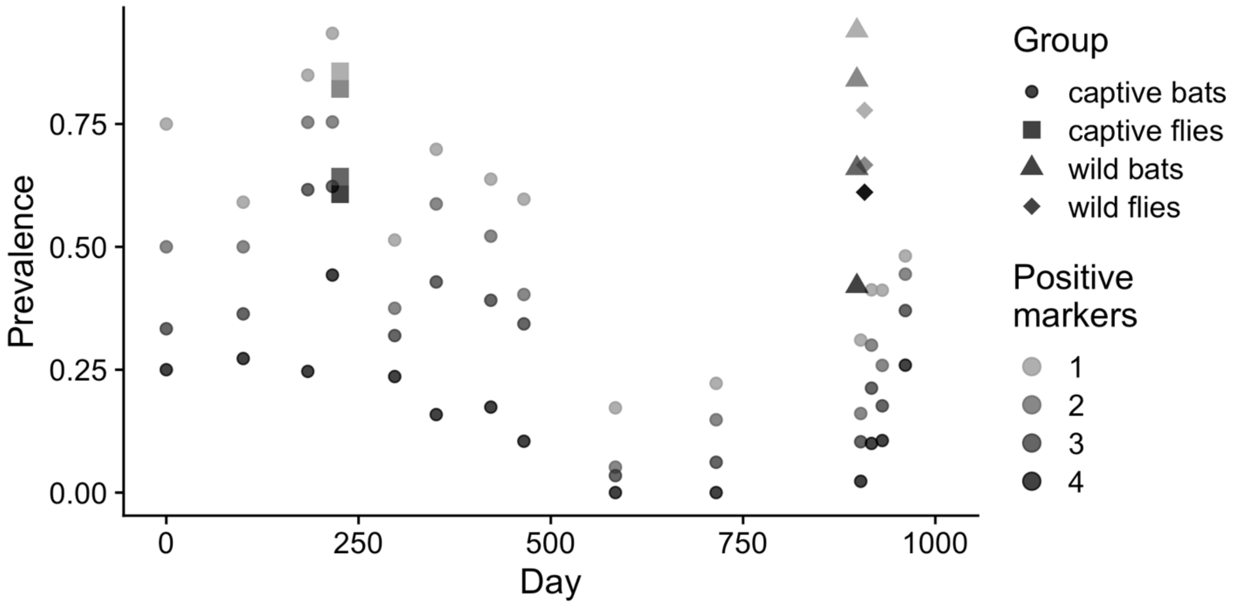


**Figure S7.** *Bartonella* infection prevalence according to the number of markers testing positive. Separate points are drawn for prevalence estimates in the *E. helvum* colony over time considering one or more, two or more, three or more, or all four markers positive (including RT-PCR). Points for sampled bat flies and wild bats are shown as unique symbols.

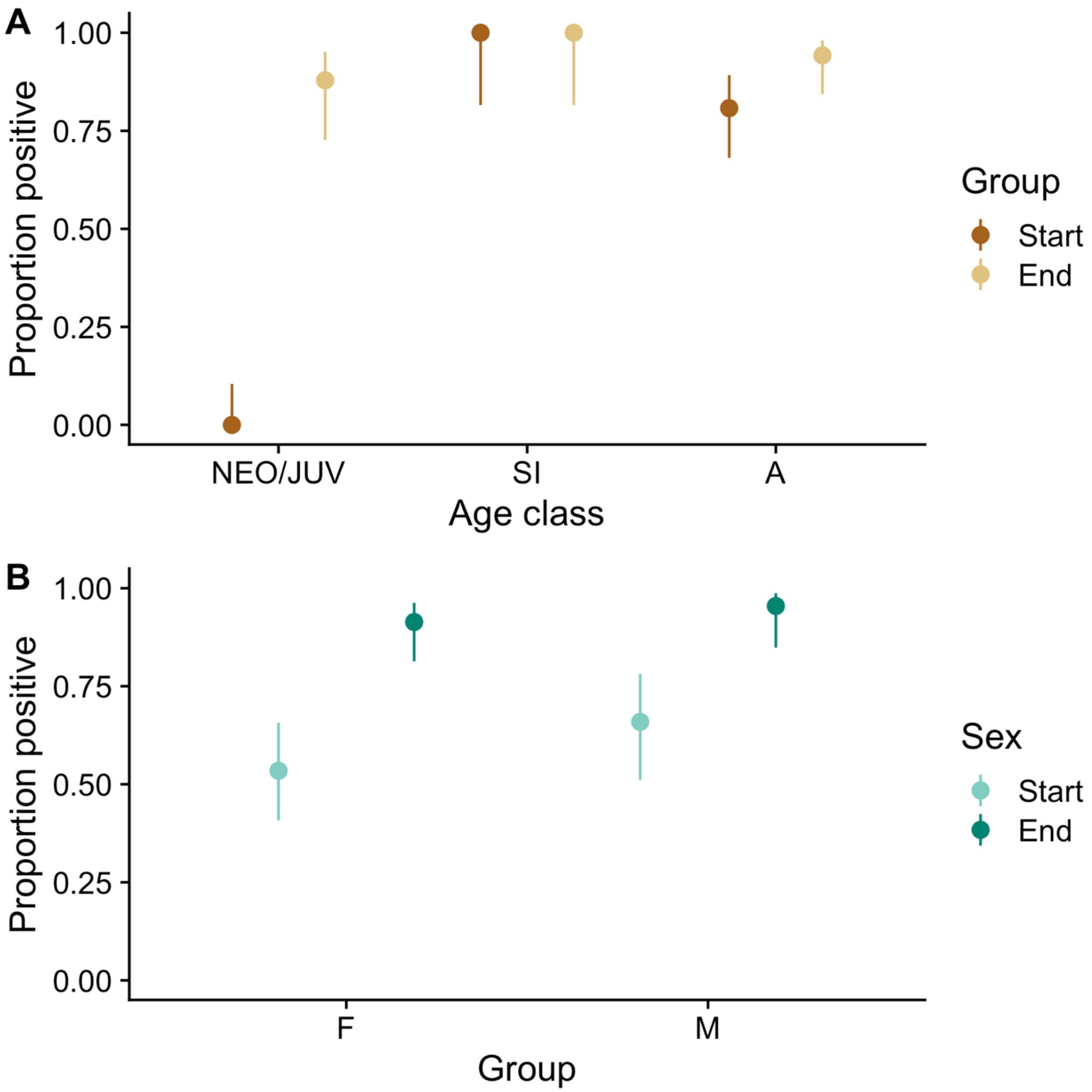


**Figure S8.** Change in the proportion of individuals positive for *Bartonella* at the start (upon entry into colony) and end (15 March 2012) of the experiment according to (A) age class and (B) sex. Wilson score 95% confidence intervals are included as lines.

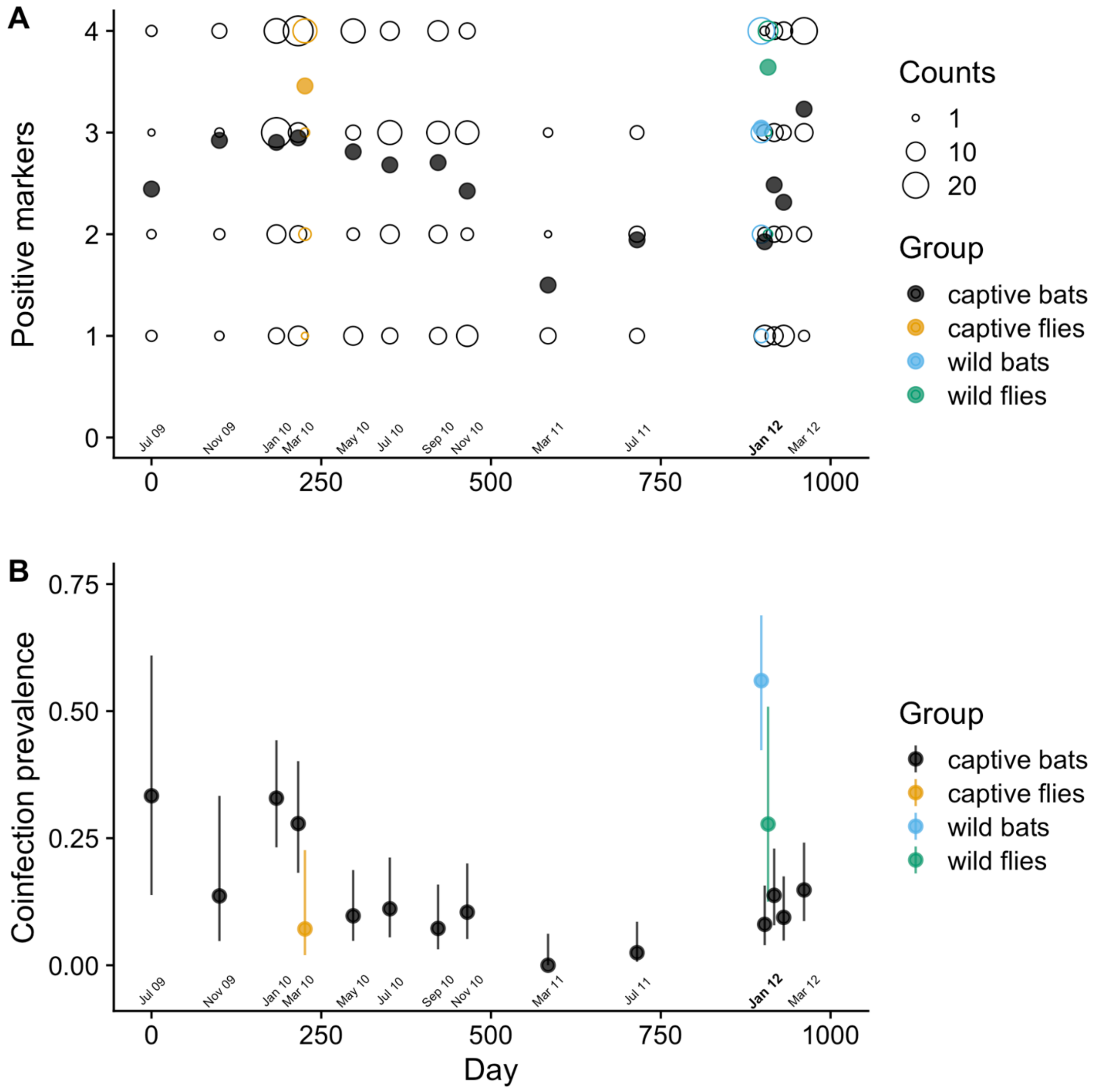


**Figure S9.** *Bartonella* infection load according to number of positive markers from each positive individual and coinfection prevalence in the *E. helvum* colony. (A) Open circles show the number of markers that were positive for each bat and bat flies with the width proportional to the number of individuals positive at that many markers. Filled circles show the mean values. (B) Coinfection prevalence was measured by the number of bats and bat flies that were positive for two or more *Bartonella* species at each time point. Wilson score 95% confidence intervals were drawn around prevalence estimates at each sampling time point.

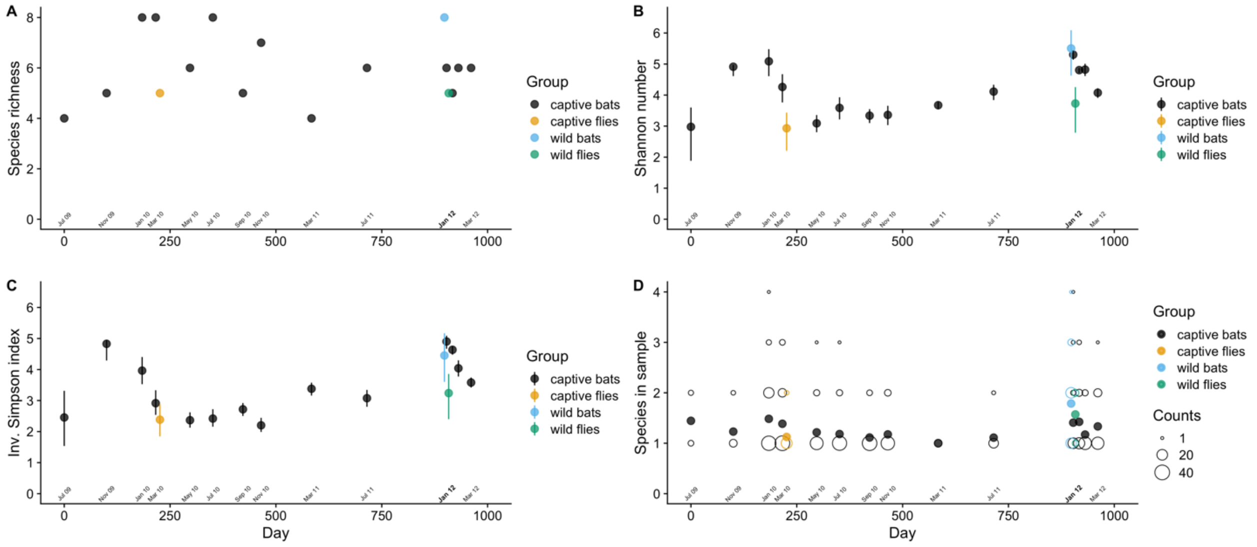


**Figure S10.** *Bartonella* infection diversity measures over time. Species richness (A), Shannon index of species evenness (B), and inverse Simpson index are colony-level measures or *Bartonella* alpha diversity. The number of *Bartonella* species in each individual bat is a measure of individual-level diversity. (B-C) Intervals around evenness indices are bootstrap 95% confidence intervals from 1000 samples from the observed multinomial distribution of *Bartonella* species relative abundances. (D) Points (open circles) show the number of *Bartonella* species observed for each individual with the width proportional to the number of individuals with that same number of *Bartonella* species. Mean values are drawn as filled circles. The month labeled in bold font on the x-axis shows when bat flies were reintroduced.

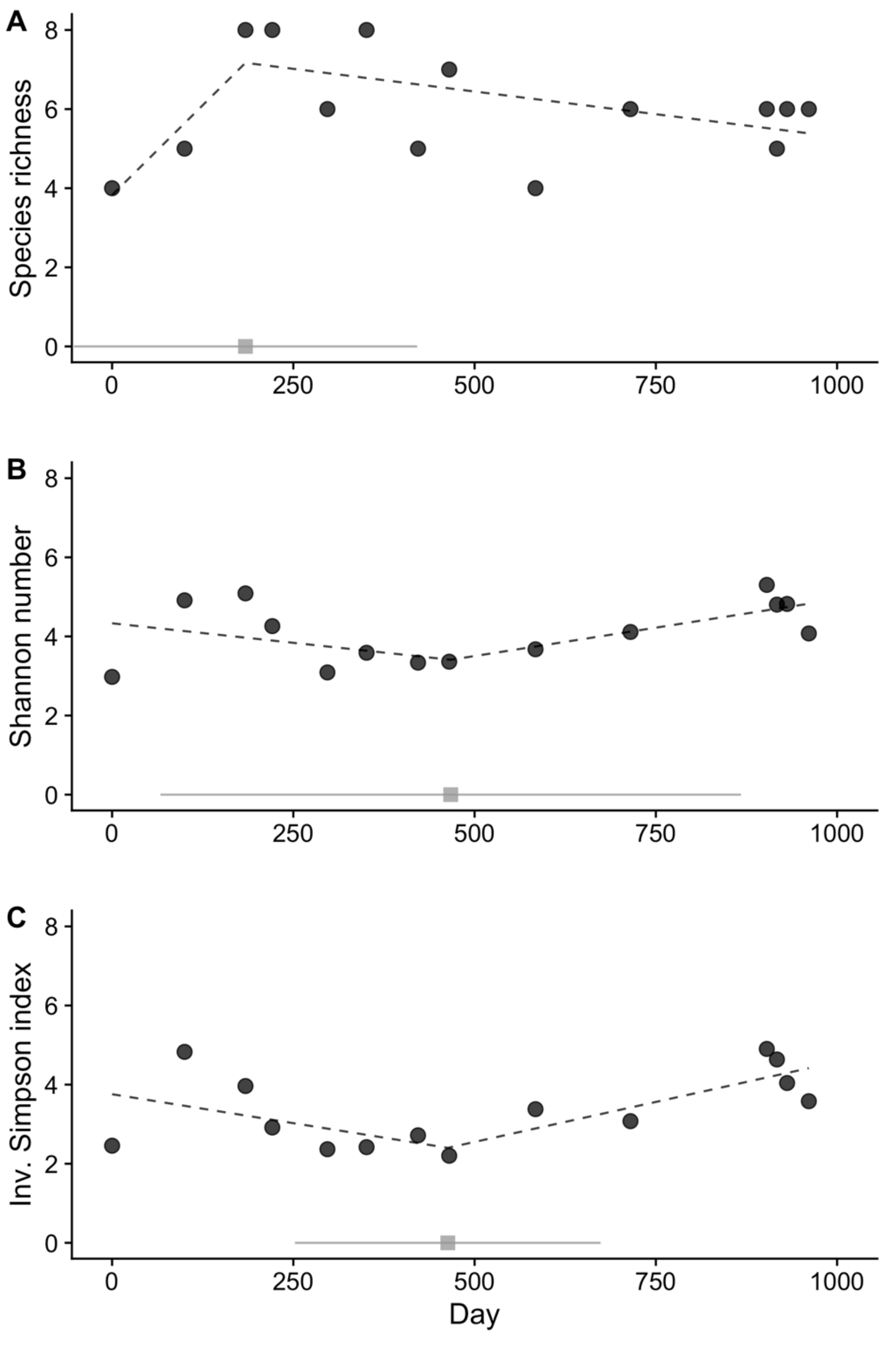


**Figure S11.** Segmented regression analysis of colony-level *Bartonella* diversity measures: (A) species richness, (B) Shannon index of species evenness, and (C) inverse Simpson index of species evenness. For each measure, dashed lines for the predicted trend from segmented regression are drawn over the data points. Breakpoints and 95% confidence intervals estimated by segmented regression are shown above the x-axis as grey squares and solid lines.

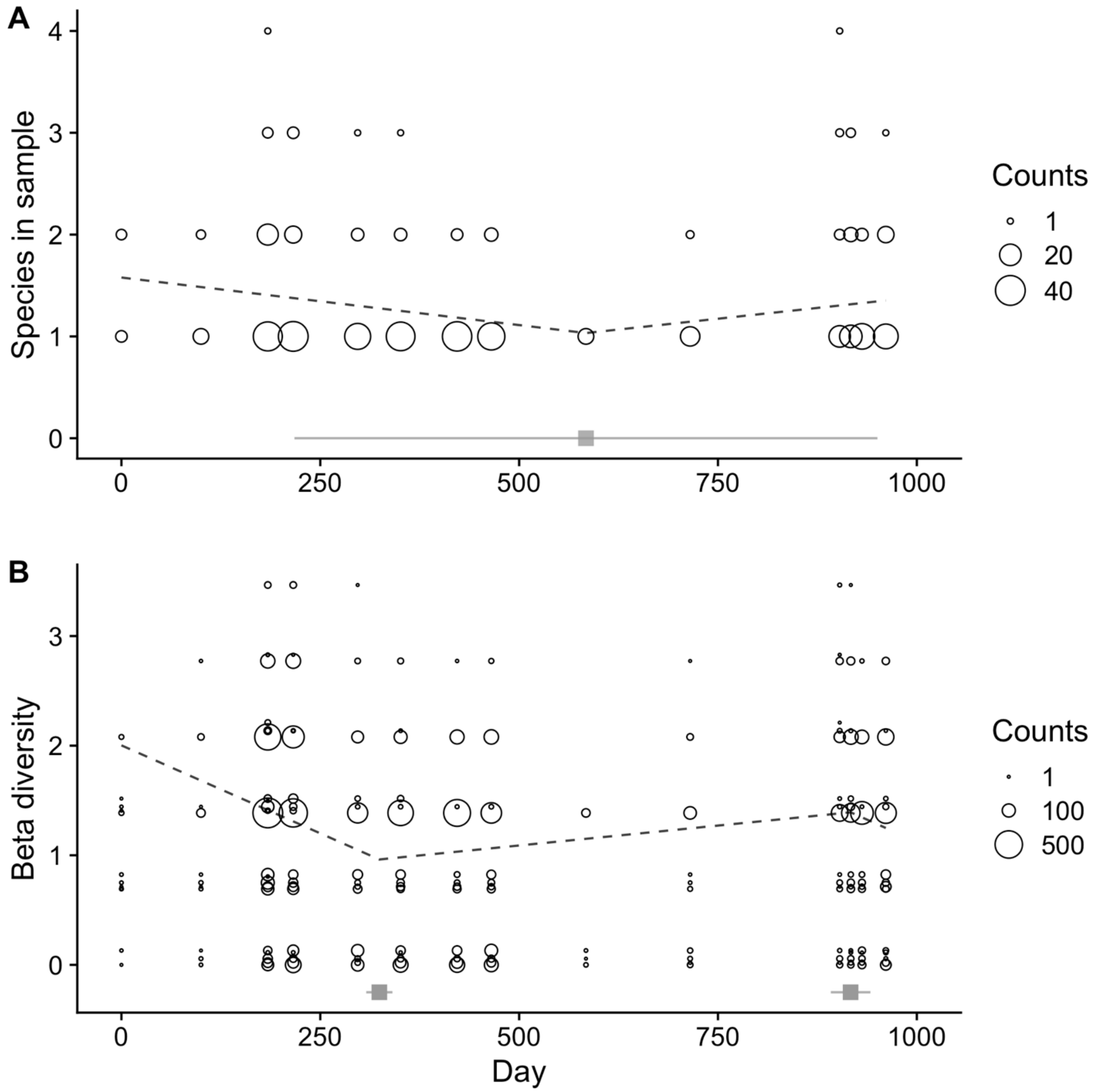


**Figure S12.** Segmented regression analysis of individual-level *Bartonella* diversity measures. Points show the number of *Bartonella* species observed in an individual sample (A) and the binomial index of beta diversity (B; compared to all other bats in the colony) for each individual with the width proportional to the number of individuals with that same diversity value. For each measure, dashed lines for the predicted trend from segmented regression are drawn over the data points. Breakpoints and 95% confidence intervals estimated by segmented regression are shown above the x-axis as grey squares and solid lines.

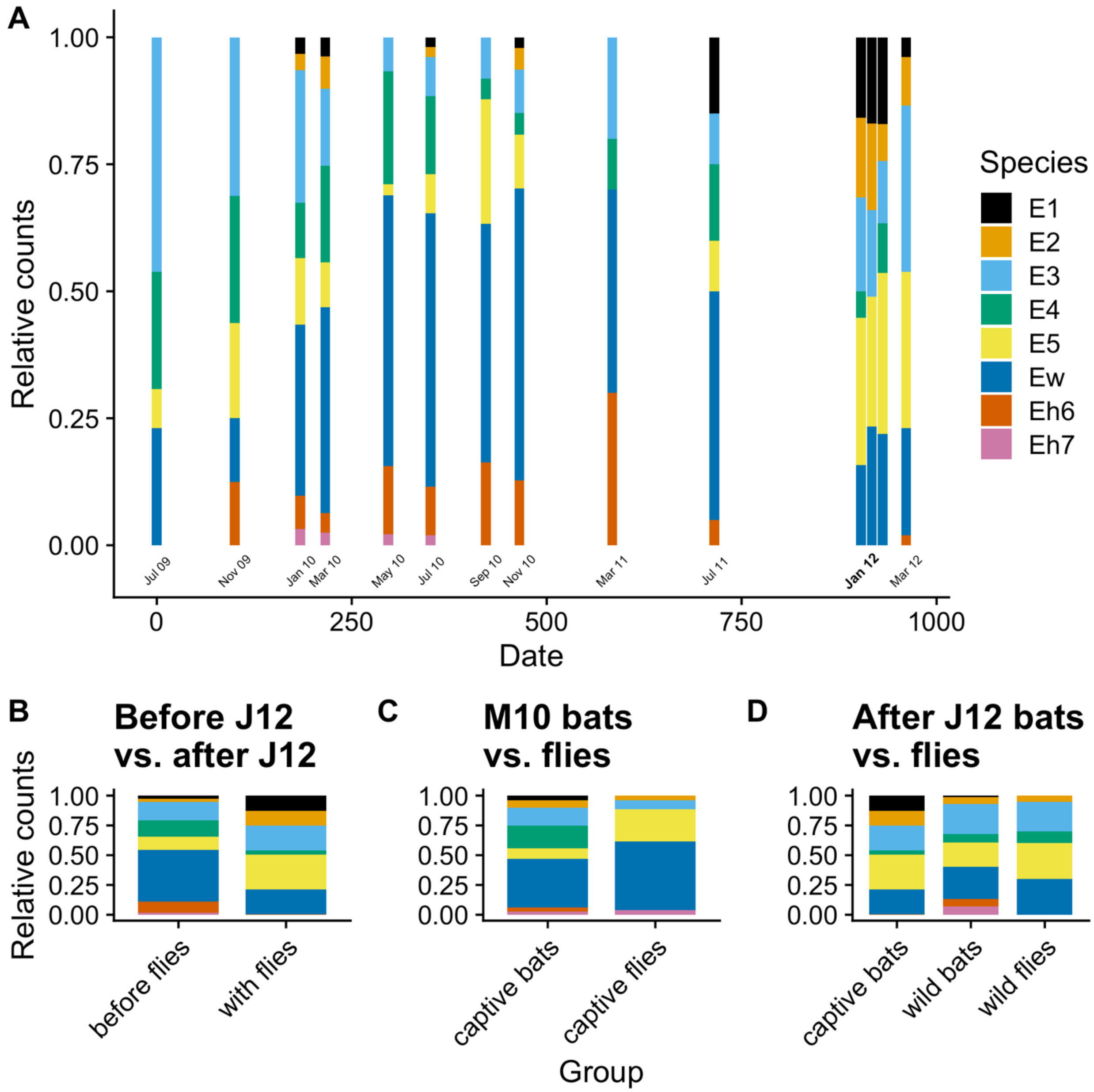


**Figure S13.** Relative counts of *Bartonella* species in the captive colony over time (A–B) and between sampled bat flies and their respective bat populations (C–D). Relative counts (A) at each time point were estimated from presence *Bartonella* species based on any positive sequence from ITS, *gltA*, and *ftsZ*. For panels A and B, the month labeled in bold font on the x-axis shows when bat flies were reintroduced. Tests for differences in the relative counts of species were performed between bats in the captive colony before and after bat flies were reintroduced on 17 January 2012 (B); between bat flies sampled from the colony and the captive bat population in March 2010 (C); and between bat flies and wild bats sampled on 17 January 2012 and the captive colony population after flies were reintroduced (D).

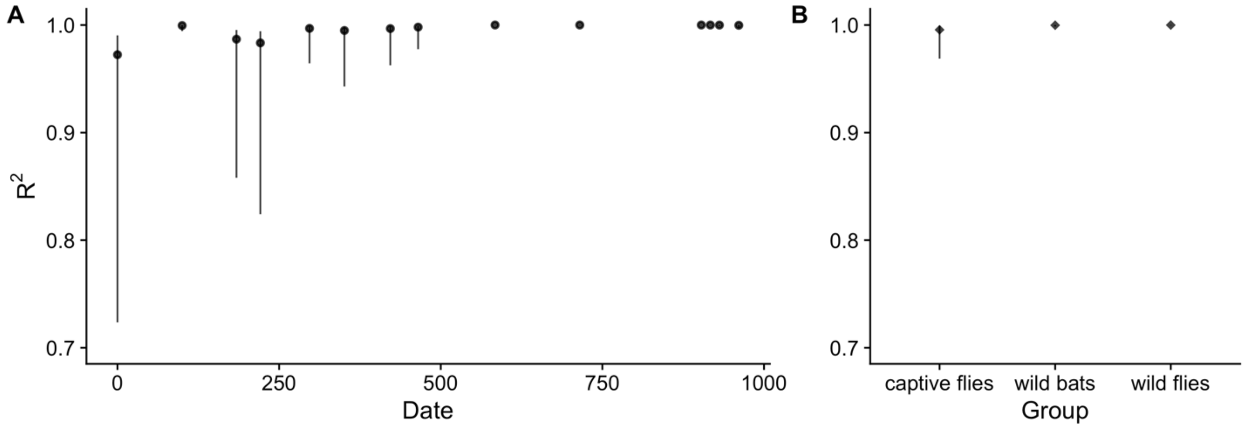


**Figure S14.** Correlation between relative abundance and relative counts of *Bartonella* species. Pearson correlation values and 95% confidence intervals are shown for each time point in the experiment (A) and for bat flies sampled on M10, wild bats sampled on J12, and wild bat flies sampled on J12 (B).

**Table S1.** Demographics of bats entering the captive colony (n = 112). Age classes follow Peel et al. (2016): neonate (NEO), juvenile (JUV), sexually immature adult (SI), and sexually mature adult (A). NEO and JUV classes were born in captivity (BIC) in 2010 and 2011 as cohorts in 4 and 5.

| | Time (days) since | | Cohort number | A | | SI | | BIC (NEO/JUV) | |
|---|---|---|---|---|---|---|---|---|---|
| Date | Study start | Last sampling | | F | M | F | M | F | M |
| 2009-07-28 | 0 | 0 | 1 | | 11 | | 1 | | |
| 2009-11-05 | 100 | 100 | 2 | 3 | 5 | 3 | 2 | | |
| 2010-01-28 | 184 | 84 | 3 | 29 | 12 | 7 | 5 | | |
| 2010-03-06 | 221 | 37 | | | | | | | |
| 2010-04-01 | 247 | No sampling | 4 | | | | | 7 | 4 |
| 2010-05-21 | 297 | 76 | | | | | | | |
| 2010-07-14 | 351 | 54 | | | | | | | |
| 2010-09-23 | 422 | 71 | | | | | | | |
| 2010-11-05 | 465 | 43 | | | | | | | |
| 2011-03-04 | 584 | 119 | | | | | | | |
| 2011-04-01 | 612 | No sampling | 5 | | | | | 14 | 9 |
| 2011-07-13 | 715 | 131 | | | | | | | |
| 2012-01-17 | 903 | 188 | | | | | | | |
| 2012-01-31 | 917 | 14 | | | | | | | |
| 2012-02-14 | 931 | 14 | | | | | | | |
| 2012-03-15 | 961 | 30 | | | | | | | |
| | | | | 32 | 28 | 10 | 8 | 21 | 13 |

**Table S2.** Oligonucleotide primers used for bacterial detection via real-time and conventional PCR amplification. Sequences designated [F] are forward primers, those designated [R] are reverse primers, and those designated [P] are TaqMan probes. FAM is 6-Carboxyfluorescein (maximum fluorescence at 518 nm).

| Bacteria | Locus | PCR type | PCR round | Primer sequence | Primer name | Reference |
|---|---|---|---|---|---|---|
| *Bartonella* | *ftsZ* | conventional | 1 | ATTAATCTGCAY-CGGCCAGA [F] | Bfp1 | (Zeaiter et al., 2002) |
| | | | 1 | ACVGADACACGA-ATAACACC [R] | Bfp2 | |
| | | conventional | 2 | ATATCGCGGAAT-TGAAGCC [F] | ftsZ R83 | (Colborn et al., 2010) |
| | | | 2 | CGCATAGAAGTA-TCATCCA [R] | ftsZ L83 | |
| *Bartonella* | *gltA* | conventional | 1 | GCTATGTCTGCA-TTCTATCA [F] | CS443f | (Birtles & Raoult, 1996; Gundi et al., 2012) |
| | | | 1 | GATCYTCAATCAT-TTCTTTCCA [R] | CS1210r | |
| | | conventional | 2 | GGGGACCAGCTC-ATGGTGG [F] | BhCS781.p | (Birtles & Raoult, 1996; Norman et al., 1995) |
| | | | 2 | AATGCAAAAAGA-ACAGTAAACA [R] | BhCS1137.n | |
| *Bartonella* | ITS | conventional | 1 | CTTCAGATGATG-ATCCCAAGCCTT-CTGGCG [F] | 325s | (Diniz et al., 2007) |
| | | | 1 | GAACCGACGACC-CCCTGCTTGCAA-AGA [R] | 1100as | |
| *Bartonella* | *ssrA* | real-time | 1 | GCTATGGTAATA-AATGGACAATGA-AATAA [F] | ssrA-F | (Diaz et al., 2012) |
| | | | 1 | GCTTCTGTTGCC-AGGTG [R] | ssrA-R | |
| | | | 1 | (FAM)-ACCCCGCTT-AAACCTGCGACG-(BHQ1) [P] | ssrA-P | |
| *Rickettsia* | 23S rRNA | real-time | 1 | AGCTTGCTTTTG-GATCATTTGG [F] | PanR8-F | (Kato et al., 2013) |
| | | | 1 | TTCCTTGCCTTT-TCATACATCTAGT [R] | PanR8-R | |
| | | | 1 | (FAM)-CCTGCTTCTATT-TGTCTTGCAGTA- | PanR8-P | |

| Bacteria | Locus | PCR type | PCR round | Primer sequence | Primer name | Reference |
|---|---|---|---|---|---|---|
| | | | | ACACGCCA-(BHQ1) [P] | | |
| *Rickettsia* | *gltA* | conventional | 1 | GGGGGCCTGCTC-ACGGCGG [F] | RpCS.877p | (Regnery et al., 1991) |
| | | | 1 | ATTGCAAAAAGT-ACAGTGAACA [R] | RpCS1258n | |
| | | conventional | 2 | GGCTAATGAAGC-AGTGATAA [F] | RpCS896p | (Choi et al., 2005; Lee et al., 2014) |
| | | | 2 | GCGACGGTATAC-CCATAGC [R] | RpCS1233n | |

**Table S3.** Thermocycler protocols used for bacterial detection via via real-time and conventional PCR amplification.

| Bacteria | Locus | PCR type | PCR round | Thermal program |
|---|---|---|---|---|
| *Bartonella* | *ftsZ* | conventional | 1 | 95°C 4:00, (95°C 0:30, 55°C 0:30, 72°C 1:00)x40, 72°C 10:00, 4°C ∞ |
| | | | 2 | 95°C 4:00, (95°C 0:30, 55°C 0:30, 72°C 1:00)x40, 72°C 10:00, 4°C ∞ |
| *Bartonella* | *gltA* | conventional | 1 | 95°C 2:00, (95°C 0:30, 48°C 0:30, 72°C 2:00)x40, 72°C 7:00, 4°C ∞ |
| | | | 2 | 95°C 3:00, (95°C 0:30, 55°C 0:30, 72°C 0:30)x40, 72°C 7:00, 4°C ∞ |
| *Bartonella* | ITS | conventional | 1 | 95°C 3:00, (95°C 0:30, 66°C 0:30, 72°C 0:30)x55, 72°C 5:00, 4°C ∞ |
| *Bartonella* | *ssrA* | real-time [1] | 1 | 60°C 1:00, 95°C 10:00, (95°C 0:15, 60°C 1:00)x45, 60°C 1:00, 4°C ∞ |
| *Rickettsia* | 23S rRNA | real-time [2] | 1 | 95°C 8:00, (95°C 0:05, 60°C 0:30)x45, 4°C ∞ |
| *Rickettsia* | *gltA* | conventional | 1 | 95°C 2:00, (95°C 0:20, 48°C 0:30, 60°C 2:00)x35, 4°C ∞ |
| | | | 2 | 95°C 10:00, (95°C 0:30, 55°C 0:30, 72°C 1:00)x30, 72°C 5:00, 4°C ∞ |

**Table S4.** Segmented regression analysis of *Bartonella* prevalence, load, and diversity. Coefficients and confidence intervals were estimated for change in slope at breakpoints. Statistical significance of parameters is indicated with an asterisk based on whether the confidence intervals overlap zero.

| Regression variable | Model family (link) | AICc | Coefficient | Estimate (95% CI) |
|---|---|---|---|---|
| Infection prevalence | Binomial (logit link) | 707.9 | Change1-2 | -0.01 (-0.014, -0.007)* |
| | | | Change2-3 | 0.025 (0.019, 0.031)* |
| | | | Point1-2 | 211.5 (178.5, 244.6) |
| | | | Point2-3 | 833.2 (806, 860.5) |
| Ct value | Gamma (inverse) | 1029.4 | Change1-2 | -0.000037 (-0.000054, -0.00002)* |
| | | | Change2-3 | 0.000056 (0.000013, 0.0001)* |
| | | | Point1-2 | 216 (171.3, 260.7) |
| | | | Point2-3 | 884.5 (819.1, 949.8) |
| Positive markers | Poisson (log) | 1540.4 | Change1-2 | -0.0017 (-0.0037, 0.00025) |
| | | | Change2-3 | 0.009 (0.0041, 0.014)* |
| | | | Point1-2 | 249.8 (118.3, 381.4) |
| | | | Point2-3 | 871.2 (815.6, 926.8) |
| Coinfection prevalence | Binomial (logit link) | 86.6 | Change1-2 | -0.0063 (-0.015, 0.0025) |
| | | | Change2-3 | 0.017 (-0.0017, 0.036) |
| | | | Point1-2 | 224.6 (38.4, 410.7) |
| | | | Point2-3 | 704.8 (455, 954.6) |
| Species richness | Poisson (log) | 65.5 | Change1-2 | -0.0038 (-0.011, 0.0038) |
| | | | Point1-2 | 184 (-52.8, 420.8) |
| Shannon number | Gamma (identity) | 43.2 | Change1-2 | 0.0048 (-0.001, 0.011) |
| | | | Point1-2 | 467.1 (66.7, 867.5) |
| Inv. Simpson index | Gamma (identity) | 44.2 | Change1-2 | 0.007 (0.0014, 0.012)* |
| | | | Point1-2 | 463 (252.3, 673.8) |

| Regression variable | Model family (link) | AICc | Coefficient | Estimate (95% CI) |
|---|---|---|---|---|
| Species in sample | Poisson (log) | 1107.9 | Change1-2 | 0.0014 (-0.00019, 0.0031) |
| | | | Point1-2 | 584 (217.6, 950.4) |
| Beta diversity | Gamma (identity) | 7764.2 | Change1-2 | 0.0039 (0.0034, 0.0045)* |
| | | | Change2-3 | -0.0039 (-0.0059, -0.002)* |
| | | | Point1-2 | 324.5 (308, 341) |
| | | | Point2-3 | 916.4 (891.4, 941.3) |

**Table S5.** Multinomial and binomial likelihood ratio (LR) tests for changes in *Bartonella* relative species abundances before and after the reintroduction of bat flies. The period before flies were reintroduced covers July 2009 to July 2011. The period after flies were reintroduced covers 17 January 2012 to March 2012.

| Date | E1 | E2 | E3 | E4 | E5 | Ew | Eh6 | Eh7 |
|---|---|---|---|---|---|---|---|---|
| 2009-07-28 | 0 | 0 | 11 | 4 | 1 | 3 | 0 | 0 |
| 2009-11-05 | 0 | 0 | 7 | 6 | 5 | 4 | 5 | 0 |
| 2010-01-28 | 3 | 4 | 37 | 16 | 19 | 60 | 7 | 4 |
| 2010-03-06 | 3 | 6 | 18 | 17 | 11 | 72 | 3 | 2 |
| 2010-05-21 | 0 | 0 | 4 | 19 | 1 | 47 | 7 | 1 |
| 2010-07-14 | 1 | 1 | 9 | 11 | 12 | 64 | 5 | 1 |
| 2010-09-23 | 0 | 0 | 7 | 4 | 33 | 54 | 8 | 0 |
| 2010-11-05 | 1 | 3 | 6 | 5 | 9 | 57 | 6 | 0 |
| 2011-03-04 | 0 | 0 | 3 | 2 | 0 | 6 | 3 | 0 |
| 2011-07-13 | 4 | 0 | 2 | 4 | 2 | 14 | 1 | 0 |
| 2012-01-17 | 8 | 9 | 10 | 2 | 15 | 6 | 0 | 0 |
| 2012-01-31 | 14 | 10 | 9 | 0 | 19 | 17 | 0 | 0 |
| 2012-02-14 | 10 | 4 | 5 | 6 | 26 | 15 | 0 | 0 |
| 2012-03-15 | 2 | 11 | 29 | 0 | 39 | 19 | 1 | 0 |
| Total abundance | 46 | 48 | 157 | 96 | 192 | 438 | 46 | 8 |
| Sum total abundance | 1031 | | | | | | | |
| Before abundance | 12 | 14 | 104 | 88 | 93 | 381 | 45 | 8 |
| Before total abundance | 745 | | | | | | | |
| Before frequency | 0.016 | 0.019 | 0.14 | 0.12 | 0.12 | 0.51 | 0.06 | 0.011 |
| After abundance | 34 | 34 | 53 | 8 | 99 | 57 | 1 | 0 |
| Expected abundance | 4.6 | 5.4 | 39.9 | 33.8 | 35.7 | 146.3 | 17.3 | 3.1 |
| After total abundance | 286 | | | | | | | |
| After frequency | 0.12 | 0.12 | 0.19 | 0.028 | 0.35 | 0.2 | 0.0035 | 0 |
| Multinomial adjusted LR | 350.1 | | | | | | | |
| Multinomial P | 0 | | | | | | | |
| Binomial adjusted LR | 78.7 | 69.8 | 4.5 | 30.5 | 91.1 | 116.5 | 27.3 | 6.1 |
| Binomial P | 0 | 1.1E-16 | 0.034 | 3.4E-08 | 0 | 0 | 1.8E-07 | 0.014 |

**Table S6.** Multinomial and binomial likelihood ratio (LR) tests of *Bartonella* species abundance changes between groups of sampled bats and bat flies: (A) tests between captive bats and captive flies in March 2010, (B) tests between wild bats and wild flies on 17 January 2012, and (C) tests between captive bats (aggregated over the period after flies were introduced) and sampled wild flies on 17 January 2012.

A

| Date | Group | E1 | E2 | E3 | E4 | E5 | Ew | Eh6 | Eh7 |
|---|---|---|---|---|---|---|---|---|---|
| 2010-03-06 | captive bats | 3 | 6 | 18 | 17 | 11 | 72 | 3 | 2 |
| 2010-03-06 | captive flies | 0 | 3 | 4 | 0 | 19 | 35 | 0 | 1 |
| Bats total abundance | | 132 | | | | | | | |
| Bats frequency | | 0.023 | 0.045 | 0.14 | 0.13 | 0.083 | 0.55 | 0.023 | 0.015 |
| Expected abundance | | 1.4 | 2.8 | 8.5 | 8 | 5.2 | 33.8 | 1.4 | 0.9 |
| Flies total abundance | | 62 | | | | | | | |
| Flies frequency | | 0 | 0.048 | 0.065 | 0 | 0.31 | 0.56 | 0 | 0.016 |
| Multinomial adjusted LR | | 43.7 | | | | | | | |
| Multinomial P | | 2.5E-07 | | | | | | | |
| Binomial adjusted LR | | 2.7 | 0.015 | 3.3 | 16.1 | 23.8 | 0.049 | 2.7 | 0.0049 |
| Binomial P | | 0.10 | 0.9 | 0.069 | 6.1E-05 | 1.1E-06 | 0.82 | 0.1 | 0.94 |

B

| Date | Group | E1 | E2 | E3 | E4 | E5 | Ew | Eh6 | Eh7 |
|---|---|---|---|---|---|---|---|---|---|
| 2012-01-17 | wild bats | 1 | 5 | 21 | 6 | 17 | 23 | 5 | 6 |
| 2012-01-17 | wild flies | 0 | 1 | 5 | 3 | 13 | 14 | 0 | 0 |
| Bats total abundance | | 84 | | | | | | | |
| Bats frequency | | 0.012 | 0.06 | 0.25 | 0.071 | 0.20 | 0.27 | 0.06 | 0.071 |
| Expected abundance | | 0.43 | 2.1 | 9 | 2.6 | 7.3 | 9.9 | 2.1 | 2.6 |
| Flies total abundance | | 36 | | | | | | | |
| Flies frequency | | 0 | 0.028 | 0.14 | 0.083 | 0.36 | 0.39 | 0 | 0 |
| Multinomial adjusted LR | | 16.7 | | | | | | | |
| Multinomial P | | 0.019 | | | | | | | |
| Binomial adjusted LR | | 0.78 | 0.73 | 2.4 | 0.067 | 4.4 | 2 | 4 | 4.9 |
| Binomial P | | 0.38 | 0.39 | 0.12 | 0.8 | 0.036 | 0.15 | 0.045 | 0.028 |

C

| Date | Group | E1 | E2 | E3 | E4 | E5 | Ew | Eh6 | Eh7 |
|---|---|---|---|---|---|---|---|---|---|
| 2012-01-17 | captive bats | 34 | 34 | 53 | 8 | 99 | 57 | 1 | 0 |
| 2012-01-17 | wild flies | 0 | 1 | 5 | 3 | 13 | 14 | 0 | 0 |

| | | | | | | | |
|---|---|---|---|---|---|---|---|
| Bats total abundance | 286 | | | | | | |
| Bats frequency | 0.12 | 0.12 | 0.19 | 0.028 | 0.35 | 0.2 | 0.0035 |
| Expected abundance | 4.3 | 4.3 | 6.7 | 1 | 12.5 | 7.2 | 0.13 |
| Flies total abundance | 36 | | | | | | |
| Flies frequency | 0 | 0.028 | 0.14 | 0.083 | 0.36 | 0.39 | 0 |
| Multinomial adjusted LR | 16.3 | | | | | | |
| Multinomial P | 0.012 | | | | | | |
| Binomial adjusted LR | 7.2 | 3.1 | 0.44 | 2.1 | 0.028 | 5.4 | 0.2 |
| Binomial P | 0.0073 | 0.076 | 0.51 | 0.15 | 0.87 | 0.02 | 0.66 |

**Table S7.** Multinomial and binomial likelihood ratio (LR) tests for changes in *Bartonella* relative species counts before and after the reintroduction of bat flies. The period before flies were reintroduced covers July 2009 to July 2011. The period after flies were reintroduced covers 17 January 2012 to March 2012.

| Date | E1 | E2 | E3 | E4 | E5 | Ew | Eh6 | Eh7 |
|---|---|---|---|---|---|---|---|---|
| 2009-07-28 | 0 | 0 | 6 | 3 | 1 | 3 | 0 | 0 |
| 2009-11-05 | 0 | 0 | 5 | 4 | 3 | 2 | 2 | 0 |
| 2010-01-28 | 3 | 3 | 24 | 10 | 12 | 31 | 6 | 3 |
| 2010-03-06 | 3 | 5 | 12 | 15 | 7 | 32 | 3 | 2 |
| 2010-05-21 | 0 | 0 | 3 | 10 | 1 | 24 | 6 | 1 |
| 2010-07-14 | 1 | 1 | 4 | 8 | 4 | 28 | 5 | 1 |
| 2010-09-23 | 0 | 0 | 4 | 2 | 12 | 23 | 8 | 0 |
| 2010-11-05 | 1 | 2 | 4 | 2 | 5 | 27 | 6 | 0 |
| 2011-03-04 | 0 | 0 | 2 | 1 | 0 | 4 | 3 | 0 |
| 2011-07-13 | 3 | 0 | 2 | 3 | 2 | 9 | 1 | 0 |
| 2012-01-17 | 6 | 6 | 7 | 2 | 11 | 6 | 0 | 0 |
| 2012-01-31 | 8 | 8 | 8 | 0 | 12 | 11 | 0 | 0 |
| 2012-02-14 | 7 | 3 | 5 | 4 | 13 | 9 | 0 | 0 |
| 2012-03-15 | 2 | 5 | 17 | 0 | 16 | 11 | 1 | 0 |
| Total counts | 34 | 33 | 103 | 64 | 99 | 220 | 41 | 7 |
| Sum total counts | 601 | | | | | | | |
| Before counts | 11 | 11 | 66 | 58 | 47 | 183 | 40 | 7 |
| Before total counts | 423 | | | | | | | |
| Before frequency | 0.026 | 0.026 | 0.16 | 0.14 | 0.11 | 0.43 | 0.095 | 0.017 |
| After counts | 23 | 22 | 37 | 6 | 52 | 37 | 1 | 0 |
| Expected counts | 4.6 | 4.6 | 27.8 | 24.4 | 19.8 | 77 | 16.8 | 2.9 |
| After total counts | 178 | | | | | | | |
| After frequency | 0.13 | 0.12 | 0.21 | 0.034 | 0.29 | 0.21 | 0.0056 | 0 |
| Multinomial adjusted LR | 183.3 | | | | | | | |
| Multinomial P | 0 | | | | | | | |
| Binomial adjusted LR | 38.2 | 34.9 | 3.3 | 21.6 | 42.2 | 39 | 26.9 | 5.8 |
| Binomial P | 6.4E-10 | 3.5E-09 | 0.07 | 3.3E-06 | 8.2E-11 | 4.2E-10 | 2.1E-07 | 0.016 |
